# A systematic analysis of rodent models implicates adipogenesis and innate immunity in pathogenesis of fatty liver disease

**DOI:** 10.1101/2020.12.15.422799

**Authors:** Yu Ri Im, Harriet Hunter, Dana de Gracia Hahn, Amedine Duret, Qinrong Cheah, Jiawen Dong, Madison Fairey, Clarissa Hjalmarsson, Alice Li, Hong Kai Lim, Lorcán McKeown, Claudia-Gabriela Mitrofan, Raunak Rao, Mrudula Utukuri, Ian A. Rowe, Jake P. Mann

## Abstract

Animal models of human disease are a key component of translational research and yet there is often no consensus on which model is optimal for a particular disease. Here, we generated a database of 3,920 rodent models of non-alcoholic fatty liver disease (NAFLD). Study designs were highly heterogeneous therefore few models had been cited more than once. Analysis of genetic models provided evidence for the role of adipose dysfunction and perturbation of the innate immune system in the progression of NAFLD. We identified that high-fat, high-fructose diets most closely recapitulate the human phenotype of NAFLD. There was substantial variability in the nomenclature of animal models; a consensus on terminology of specialist diets is needed. More broadly, this analysis demonstrates the variability in preclinical study design, which has implications for the reproducibility of *in vivo* experiments.

## Introduction

Animal models of human diseases are an integral part of pre-clinical research in order to facilitate mechanistic or therapeutic studies that are not possible in humans. Studies of non-alcoholic fatty liver disease (NAFLD) predominantly utilise mostly rodent models of obesity (or insulin resistance) to induce hepatic steatosis(Friedman et al., 2018; Hebbard and George, 2011). However, it is not clear which study design most closely reflects the human disease phenotype.

NAFLD is a slowly progressive condition characterised by accumulation of excess hepatic lipids(Diehl and Day, 2017). Some individuals develop non-alcoholic steatohepatitis (NASH) that drives fibrosis, which can lead to cirrhosis and hepatocellular carcinoma (HCC). NAFLD is causally associated with insulin resistance(Dongiovanni et al., 2018; Liu et al., 2020), typically via obesity, which has been reflected in the recently coined terminology ‘metabolic-dysfunction associated fatty liver disease’ (Eslam et al., 2020). It is also recognised that individuals with common variants that exacerbate NAFLD (e.g. p.Ile148Met in *PNPLA3*) may develop NAFLD despite being less insulin resistant(Luukkonen et al., 2016). Almost all such variants perturb hepatic lipid metabolism(Mann et al., 2020), whereas there has been little human genetic evidence for the inflammatory or fibrotic aspects of NASH.

The theoretical ‘ideal’ NAFLD model should reflect the full human spectrum of hepatic disease plus features of the associated metabolic syndrome and would pass through these stages without taking an unacceptably long duration (e.g. >1 year)(Castro and Diehl, 2018; Febbraio et al., 2019). Whilst there have been many narrative reviews on the topic, there has not previously been a systematic categorisation and assessment of animal models.

We have recently found that variation in study design (e.g. %kcal fat in diet, age of mice) affects the treatment response in rodent models of NAFLD(Hunter et al., 2020). Therefore, detailed appraisal of pre-clinical study design is important for reproducibility, particularly given recent high-profile studies that could not be replicated.

In order to address these questions, we systematically reviewed and categorised the NAFLD models from over 4500 published studies. We then used data from genetically modified animals to identify pathways involved with development of NAFLD.

## Results

### The landscape of animal models of NAFLD

In order to understand the spectrum of animal models used to study fatty liver disease, we performed a systematic search of multiple databases for all published papers related to NAFLD, NASH, and hepatic steatosis. 8727 articles were screened and 4540 articles were ultimately included in our analysis (Figure 1).

**Fig. 1:**
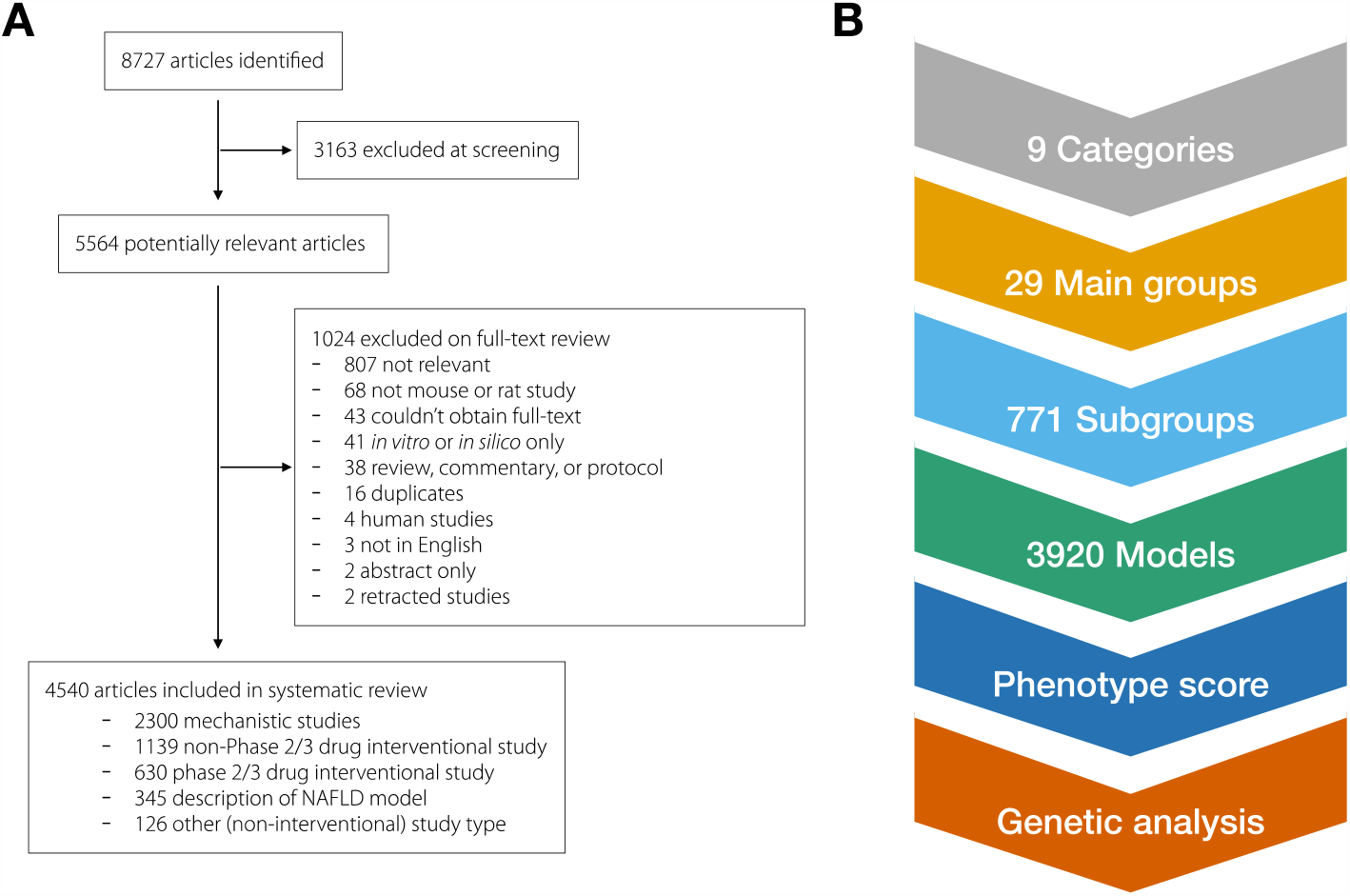
Study design. (A) Article inclusion and exclusion flow chart. (B) Overview of the categorisation hierarchy used in analysis.

From this, we built a database of 3920 unique rodent models of fatty liver disease, which we grouped into nine broad categories of model (Table 1). These could be further subdivided into 29 Main Groups (E.g. “High fat, high cholesterol diet”) and 771 Subgroups (e.g. “ApoE mutant with dietary manipulation”). Most models were dietary (1927/3920), followed by genetically altered rodents combined with dietary manipulation (826/3920).

**Table 1:** Characteristics of rodent models of NAFLD. Data from 4540 studies reporting 3920 unique NAFLD models, divided into nine categories. SE, standard error.

| Category |  | Overall | Dietary | Dietary + other | Genetic | Genetic + chemical | Genetic + dietary | Genetic + other | Chemical | Offspring | Other |
| --- | --- | --- | --- | --- | --- | --- | --- | --- | --- | --- | --- |
| Number of models |  | 3920 | 1927 | 68 | 477 | 65 | 826 | 26 | 391 | 91 | 49 |
| Main groups |  | 29 | 14 | 9 | 8 | 7 | 8 | 6 | 3 | 4 | 4 |
| Subgroups |  | 771 | 44 | 16 | 275 | 32 | 359 | 23 | 28 | 15 | 14 |
| Sex | Both | 175 (5.2%) | 81 (4.7%) | 2 (3%) | 14 (4.3%) |  | 47 (6.7%) | 1 (4.8%) | 10 (2.7%) | 19 (22.1%) | 1 (2.3%) |
|  | Female | 444 (13.1%) | 180 (10.4%) | 32 (48.5%) | 47 (14.6%) | 7 (13.7%) | 78 (11.1%) | 5 (23.8%) | 52 (14.2%) | 26 (30.2%) | 17 (38.6%) |
|  | Male | 2777 (81.8%) | 1476 (85%) | 32 (48.5%) | 261 (81.1%) | 44 (86.3%) | 578 (82.2%) | 15 (71.4%) | 304 (83.1%) | 41 (47.7%) | 26 (59.1%) |
| Age diet started | Median (SE) | 7.0 (0.1) | 7.0 (0.1) | 7.8 (0.3) |  | 8.0 (0.8) | 8.0 (0.2) | 8.0 (0.8) | 6.0 (0.2) | 4.0 (0.5) | 8.0 (0.6) |
|  | Range | (1.0 - 101.0) | (1.0 - 101.0) | (2.0 - 14.0) |  | (3.0 - 36.0) | (2.0 - 62.0) | (4.0 - 16.0) | (3.0 - 32.0) | (3.0 - 21.0) | (3.0 - 21.0) |
| Fat in diet (%kcal) | Median (SE) | 45.0 (0.3) | 44.5 (0.4) | 45.0 (1.8) |  | 45.0 (1.8) | 45.0 (0.5) | 42.5 (2.9) | 53.0 (0.9) | 43.0 (1.5) | 34.7 (3.1) |
|  | Range | (0.0 - 100.0) | (0.0 - 95.1) | (0.1 - 82.0) |  | (12.0 - 60.0) | (0.0 - 100.0) | (16.7 - 60.0) | (5.0 - 82.0) | (14.2 - 60.0) | (11.1 - 60.0) |
| Cholesterol in diet (%weight) | Median (SE) | 1.0 (0.03) | 1.2 (0.05) | 0.5 (0.30) |  | 0.5 (0.06) | 0.2 (0.05) |  | 2.0 (0.14) | 0.3 (0.07) | 2.0 (0.23) |
|  | Range | (0.0 - 15.0) | (0.0 - 15.0) | (0.0 - 10.0) |  | (0.2 - 1.2) | (0.0 - 12.5) |  | (0.0 - 10.0) | (0.1 - 2.0) | (0.0 - 4.0) |
| Sucrose in diet (%kcal) | Median (SE) | 29.0 (0.3) | 26.2 (0.4) | 32.0 (1.8) |  | 17.4 (1.5) | 30.4 (0.6) |  | 18.2 (1.3) | 30.2 (1.9) |  |
|  | Range | (1.0 - 88.9) | (1.0 - 88.9) | (21.7 - 42.4) |  | (10.3 - 34.0) | (6.7 - 74.0) |  | (6.5 - 73.0) | (27.0 - 65.2) |  |
| Fructose in diet (%weight) | Median (SE) | 21.0 (0.3) | 20.0 (0.5) | 13.8 (1.4) |  |  | 30.0 (0.8) |  | 26.0 (0.9) | 20.0 (0.0) | 37.5 (4.5) |
|  | Range | (2.3 - 79.0) | (2.3 - 79.0) | (2.3 - 30.0) |  |  | (2.3 - 70.0) |  | (2.3 - 66.0) | (20.0 - 20.0) | (15.0 - 60.0) |
| Lipodystrophy |  | 48 (1.2%) |  |  | 33 ( 6.9%) |  | 15 ( 1.8%) |  |  |  |  |
| Metabolic syndrome features | Obesity | 1855 (47.3%) | 1011 (52.5%) | 41 (60.3%) | 154 (32.3%) | 31 (47.7%) | 440 (53.3%) | 10 (38.5%) | 100 (25.6%) | 47 (51.6%) | 21 (42.9%) |
|  | Insulin resistant | 1709 (43.6%) | 846 (43.9%) | 24 (35.3%) | 212 (44.4%) | 23 (35.4%) | 408 (49.4%) | 11 (42.3%) | 132 (33.8%) | 36 (39.6%) | 17 (34.7%) |
|  | Dyslipidaemia | 1993 (50.8%) | 1025 (53.2%) | 27 (39.7%) | 234 (49.1%) | 35 (53.8%) | 442 (53.5%) | 15 (57.7%) | 146 (37.3%) | 46 (50.5%) | 23 (46.9%) |
| Liver histology features | Raised transaminases | 1663 (42.4%) | 880 (45.7%) | 30 (44.1%) | 139 (29.1%) | 33 (50.8%) | 350 (42.4%) | 9 (34.6%) | 177 (45.3%) | 27 (29.7%) | 18 (36.7%) |
|  | Lobular inflammation | 772 (19.7%) | 408 (21.2%) | 16 (23.5%) | 58 (12.2%) | 12 (18.5%) | 165 (20.0%) | 5 (19.2%) | 93 (23.8%) | 7 (7.7%) | 8 (16.3%) |
|  | Hepatocellular ballooning | 745 (19.0%) | 411 (21.3%) | 10 (14.7%) | 48 (10.1%) | 11 (16.9%) | 146 (17.7%) | 3 (11.5%) | 91 (23.3%) | 16 (17.6%) | 9 (18.4%) |
|  | Steatohepatitis | 1369 (34.9%) | 718 (37.3%) | 19 (27.9%) | 100 (21.0%) | 26 (40.0%) | 323 (39.1%) | 5 (19.2%) | 144 (36.8%) | 21 (23.1%) | 13 (26.5%) |
|  | (Peri-)Portal inflammation | 167 (4.3%) | 106 (5.5%) | 4 (5.9%) | 18 (3.8%) |  | 21 (2.5%) |  | 15 (3.8%) | 1 (1.1%) | 2 (4.1%) |
|  | Fibrosis | 891 (22.7%) | 455 (23.6%) | 17 (25.0%) | 75 (15.7%) | 12 (18.5%) | 215 (26.0%) | 4 (15.4%) | 97 (24.8%) | 8 (8.8%) | 8 (16.3%) |
|  | Hepatocellular carcinoma | 177 (4.5%) | 34 ( 1.8%) | 4 ( 5.9%) | 50 (10.5%) | 15 (23.1%) | 19 ( 2.3%) | 4 (15.4%) | 50 (12.8%) |  | 1 ( 2.0%) |
| Citations per model | Mean | 1.59 | 1.81 | 1.04 | 2.16 | 1.03 | 1.09 | 1 | 1.19 | 1.03 | 1.24 |
|  | Range | (1.0 - 90.0) | (1.0 - 78.0) | (1.0 - 2.0) | (1.0 - 90.0) | (1.0 - 2.0) | (1.0 - 6.0) | (1.0 - 1.0) | (1.0 - 12.0) | (1.0 - 2.0) | (1.0 - 4.0) |

The majority of models were conducted using only male animals (2777/3920, 82%) and 27% were performed using C57BL/6J mice, with hundreds of other genetic backgrounds represented (SupTable 1). 12-weeks was the modal duration of model study (mean = 15 weeks, Figure 2A), this was similar for all histological features except presence of HCC, which was reported at up to two years of age (SupFig 2).

**Fig. 2:**
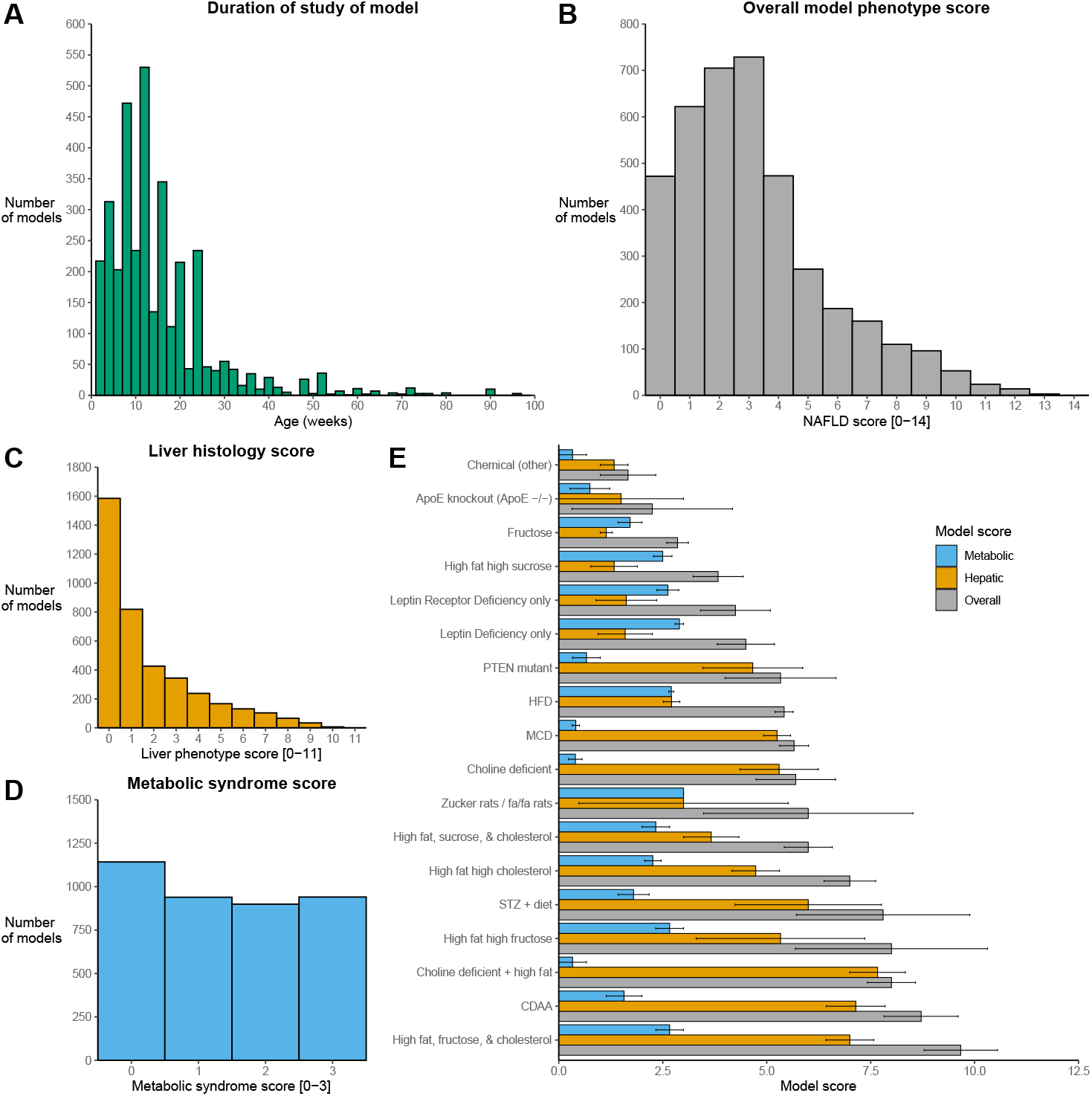
Phenotype score and duration of rodent models of NAFLD. (A) Histogram illustrating the maximum duration each model (n=3920) was studied. (B) Overall phenotype score for each model [0-14] generated from a combination of liver histology score and metabolic syndrome score. (C) Liver histology score [0-11] for each model, where one point is given for the presence of each histological feature of human NAFLD, including, fibrosis stages 1-4 and HCC, plus raised aminotransferases. (D) Metabolic syndrome score (0-3) for each model was calculated as the presence of obesity, insulin resistance, and dyslipidemia, with one point for each. (E) Comparison of mean (± standard error) scores for subgroups of models with at least 3 models, which had all been replicated at least once. The dotted line represents the mean value. ApoE, apolipoprotein E; CDAA, choline-deficient amino acid-defined diet; HFD, high fat diet; PTEN, phosphatase and tensin homolog; STZ, streptozocin.

### Replication of model design

There is an increasing awareness of the need for replication and reproducibility across all fields of preclinical research(von Herrath et al., 2019). Therefore we examined how frequently each model was used by independent studies. Due to differences in model design (e.g. proportion of fat in diet, age diet initiated, genetic background), models were used by a mean of 1.6 studies (median 1 study). Genetic models fed standard chow were used by a mean 2.2 studies, compared to only 1.1 for genetic models with dietary manipulation. There was a trend that more complex models (e.g. Offspring, Chemical plus dietary) were more likely to be used by only a single study. The most frequently used model was male leptin deficient (*ob/ob*) mice on a pure C57BL/6J background fed standard chow (Table 2).

**Table 2:**
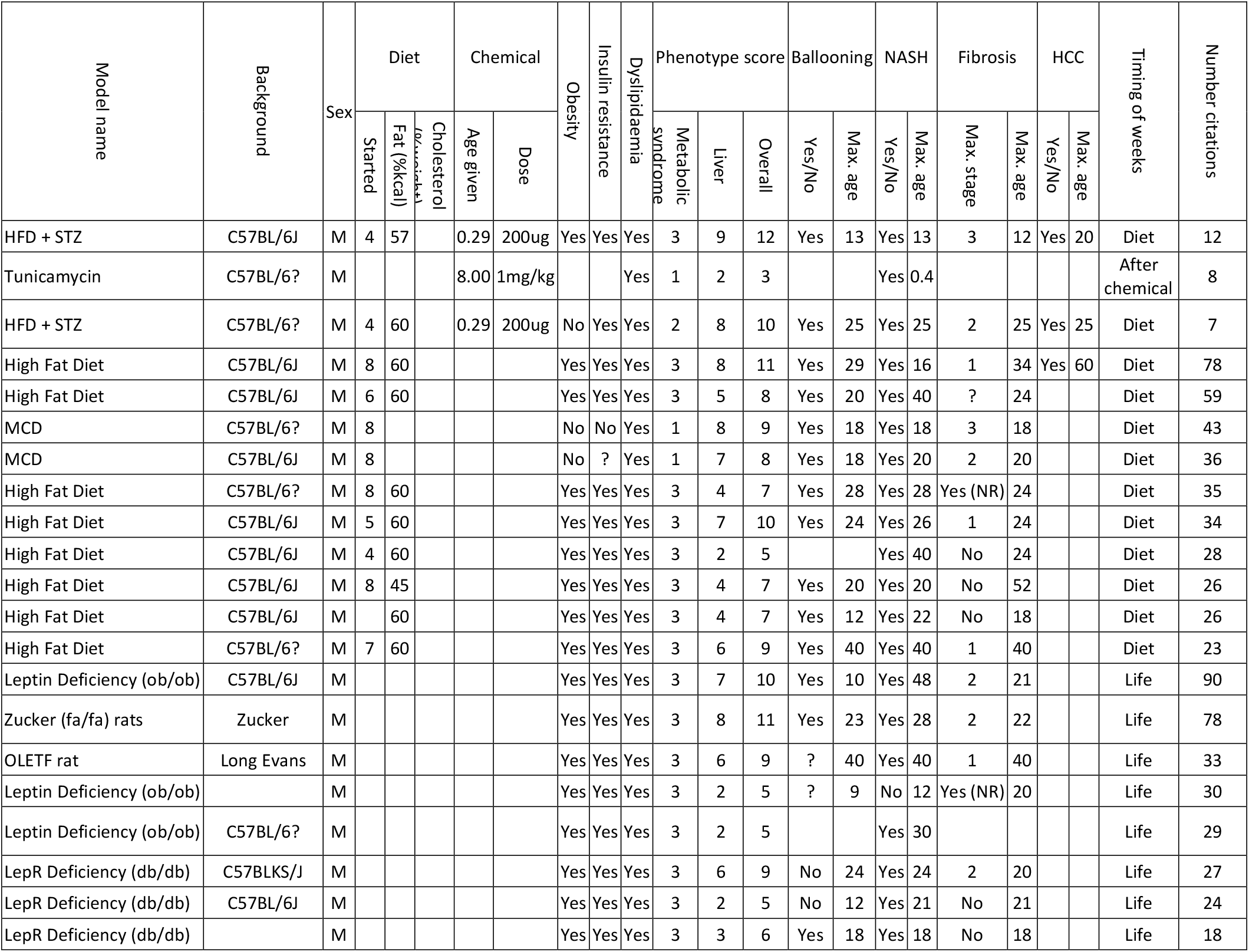
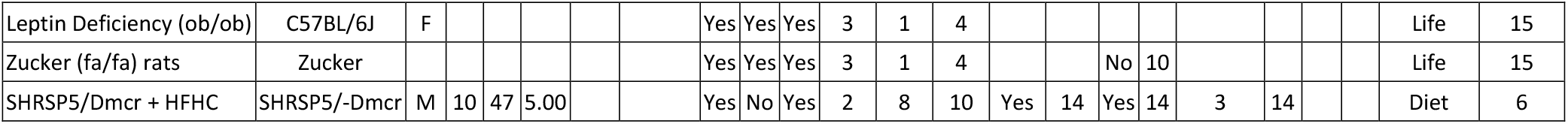
Most frequently used rodent NAFLD models. Within each of nine categories of NAFLD models, the most frequently used models (with minimum 3 citations) are reported. All ages are given in weeks. “?” is used to illustrate where there was no consensus between studies for presence of metabolic syndrome or histological features. HFD, high fat diet; HFHC, high fat high cholesterol; MCD, methionine & choline-deficient diet; OLETF, Otsuka Long-Evans Tokushima Fatty; NR, (fibrosis stage) not reported; STZ, streptozocin.

### Comparability of models to the metabolic human phenotype of NAFLD

In humans, fatty liver disease almost always occurs in concert with other features of the metabolic syndrome, driven by insulin resistance(Samuel and Shulman, 2018). This has been reflected in the recent revision of terminology that also acknowledges that individuals with the most severe insulin resistance and metabolic risk factors are also at greatest risk of progressive liver disease(Eslam et al., 2020). Therefore, we assessed whether rodent models were reported to show obesity, insulin resistance, or dyslipidemia, summarised as a ‘metabolic syndrome score (0-3)’. Overall, 44% models were reported to show insulin resistance, 47% were obese, and 51% had dyslipidemia. 47% (1839/3920) models had at least two features of the metabolic syndrome (Figure 2D). Chemically-induced models were less frequently reported to show metabolic features of NAFLD (e.g. 26% with obesity).

Fatty liver disease manifests a spectrum of histological features on liver biopsy in humans, including steatosis, lobular and periportal inflammation, hepatocellular ballooning, and fibrosis(Brunt et al., 2020). Simple steatosis (non-alcoholic fatty liver) can be distinguished from NASH through assessment by pathologists, as the latter typically requires the presence of lobular inflammation and hepatocellular ballooning. Therefore, we assessed whether each model was reported to demonstrate these features. 93% (3655/3920) models reported liver histology. Overall, 35% models were reported to show NASH and 23% to show some stage of fibrosis, whereas the presence or absence of periportal inflammation was described in only 9% of models (SupFig 1).

No model was reported to show all possible features of NAFLD liver histology and the metabolic syndrome. Three models had an overall NAFLD model score of 13 (out of 14), two of which were based on high fat, high fructose, and added cholesterol diets, though only one had been used in at least two studies (Table 3). This was supported on analysis of phenotype scores by subgroups (Figure 2E). Whilst choline-deficient diets had the highest mean liver histology scores, they had few features of the metabolic syndrome.

**Table 3:** Models with highest overall phenotype scores. Rodent models with the highest overall phenotype scores [0-14], which had been used by at least two studies. HFD, high fat diet; HFHC, high fat high cholesterol; MCD, methionine & choline-deficient diet; OLETF, Otsuka Long-Evans Tokushima Fatty; STZ, streptozocin.

| Model name | Background | Sex | Diet |  |  |  |  | Chemical |  | Obesity | Insulin resistance | Dyslipidaemia | Phenotype score |  |  | Ballooning |  | NASH |  | Fibrosis |  | HCC |  |
| --- | --- | --- | --- | --- | --- | --- | --- | --- | --- | --- | --- | --- | --- | --- | --- | --- | --- | --- | --- | --- | --- | --- | --- |
|  |  |  | Started | Fat (%kcal) | Cholesterol (%weight) | Sucrose (%kcal) | Fructose (%weight) | Age given | Dose |  |  |  | Metabolic | Liver | Overall | Yes/No | Max. age | Yes/No | Max. age | Max. stage | Max. age | Yes/No | Max. age |
| HFD + STZ | C57BL/6J | M | 4 | 57 |  |  |  | 0.3 | 200µg | Yes | Yes | Yes | 3 | 9 | 12 | Yes | 13 | Yes | 13 | 3 | 12 | Yes | 20 |
| HFD + STZ | C57BL/6J | M | 4 | 60 |  |  |  | 0.3 | 200µg | No | Yes | Yes | 2 | 9 | 11 | Yes | 5 | Yes | 20 | 3 | 20 | Yes | 20 |
| High fat, Carbohydrate, Cholesterol + Fructose in drinking water | C57BL/6J-129/SvImJ | M | 10 | 42 | 0.1 |  | 2.31 |  |  | Yes | Yes | Yes | 3 | 10 | 13 | Yes | 8 | Yes | 24 | 3 | 52 | Yes | 52 |
| High Fat Diet | C57BL/6? | M | 8 | 57 |  |  |  |  |  | Yes | Yes | Yes | 3 | 9 | 12 |  |  | Yes | 60 | 3 | 60 | Yes | 60 |
| High Fat Diet | Wistar | M | 5 | 60 |  |  |  |  |  | Yes | Yes | Yes | 3 | 9 | 12 | Yes | 8 | Yes | 8 | 3 | 8 |  |  |
| High Fat Diet | Sprague Dawley | M | 5 | 22 | 2 |  |  |  |  | Yes | Yes | Yes | 3 | 9 | 12 | Yes | 20 | Yes | 20 | 3 | 20 |  |  |
| Atherogenic + HFD | C57BL/6J | M | 8 | 60 | 1.25 |  |  |  |  | No | Yes | Yes | 2 | 10 | 12 | Yes | 68 | Yes | 68 | 4 | 68 | Yes | 68 |
| High Fat Diet | Sprague Dawley | M | 8 | 40 | 2 |  |  |  |  | Yes | Yes | Yes | 3 | 9 | 12 | Yes | 20 | Yes | 32 | 3 | 32 |  |  |
| Western diet (high fat, high fructose) | C57BL/6J | M | 8 | 58 |  |  | 2.31 |  |  | Yes | Yes | Yes | 3 | 9 | 12 | Yes | 52 | Yes | 52 | 2 | 52 | Yes | 52 |
| Choline-deficient L-amino-defined diet | Wistar | M | 6 |  |  |  |  |  |  | Yes | Yes | Yes | 3 | 8 | 11 | Yes | 4 | Yes | 12 | 3 | 12 |  |  |
| Choline-deficient L-amino-defined diet | C57BL/6? | M | 8 |  |  |  |  |  |  | No | Yes | Yes | 2 | 9 | 11 | Yes | 22 | Yes | 22 | 4 | 22 |  |  |
| Choline deficient diet | Wistar | M | 7 |  |  |  |  |  |  | No |  | Yes | 1 | 10 | 11 | Yes | 12 | Yes | 12 | 4 | 16 | Yes | 16 |
| High Fat Diet / HFD | C57BL/6J | M | 8 | 60 |  |  |  |  |  | Yes | Yes | Yes | 3 | 8 | 11 | Yes | 29 | Yes | 16 | 1 | 34 | Yes | 60 |
| High fat (trans-fat) + fructose + cholesterol | C57BL/6J | M | 7 | 40 | 2 |  | 22 |  |  | Yes | Yes | Yes | 3 | 8 | 11 | Yes | 30 | Yes | 52 | 2 | 52 |  |  |
| High fat (trans-fat) + fructose + cholesterol | C57BL/6J | M | 5 | 40 | 2 |  | 20 |  |  | Yes | Yes | Yes | 3 | 8 | 11 | Yes | 50 | Yes | 50 | 3 | 50 |  |  |
| HCHF diet + Fructose and Sucrose in water | C57BL/6J | M | 8 | 58 |  |  | 2.31 |  |  | Yes | Yes | Yes | 3 | 8 | 11 | Yes | 16 | Yes | 16 | 2 | 16 |  |  |
| Methionine and choline deficient diet / MCD | Wistar | M | 7 |  |  |  |  |  |  | No |  | Yes | 1 | 10 | 11 | Yes | 4 | Yes | 20 | 3 | 20 | Yes | 20 |
| MCD + HFD | C57BL/6? | M | 8 | 60 |  |  |  |  |  | No | Yes | Yes | 2 | 9 | 11 | Yes | 2 | Yes | 12 | 4 | 16 |  |  |
| ACOX1-Lampe1 (splice-site mutant) | C57BL/6J |  |  |  |  |  |  |  |  | Yes |  | Yes | 2 | 9 | 11 |  |  | Yes | 59 | 4 | 59 | Yes | 59 |
| Zucker rats / fa/fa rats | Zucker | M |  |  |  |  |  |  |  | Yes | Yes | Yes | 3 | 8 | 11 | Yes | 23 | Yes | 28 | 2 | 22 |  |  |
| MC4R-KO + Western diet | C57BL/6J | M | 8 | 41 | 0.21 | 34 |  |  |  | Yes | Yes | Yes | 3 | 9 | 12 | Yes | 52 | Yes | 52 | 3 | 52 | Yes | 52 |
| SHRSP5/Dmcr rats + HFHC diet | SHRSP5/Dmcr | M | 6 | 47 | 5 |  |  |  |  | Yes | Yes | Yes | 3 | 9 | 12 | Yes | 16 | Yes | 16 | 4 | 16 |  |  |
| ACOX1-Lampe1 (splice-site mutant) + HFHC diet | C57BL/6J |  |  | 58 |  |  | 2.31 |  |  | Yes |  | Yes | 2 | 9 | 11 |  |  | Yes | 52 | 4 | 52 | Yes | 59 |
| ApoE*3Leiden+ Western diet | C57BL/6J | M | 9 | 33 | 1 |  |  |  |  | Yes | Yes | Yes | 3 | 8 | 11 | Yes | 16 | Yes | 16 | 3 | 16 |  |  |
| Leptin Deficiency (ob/ob) + High fat (trans-fat), fructose, cholesterol diet | B6.V-Lepob/JRj | M | 5 | 40 | 2 |  | 20 |  |  | Yes | Yes | Yes | 3 | 8 | 11 | Yes | 20 | Yes | 20 | 3 | 30 |  |  |
| Leptin Deficiency (ob/ob) + High fat (trans-fat), fructose, cholesterol diet | C57BL/6J | M | 8 | 40 | 2 |  | 22 |  |  | Yes | Yes | Yes | 3 | 8 | 11 | Yes | 12 | Yes | 12 | 3 | 12 |  |  |

### Identification of models with rapid progression to cirrhosis or HCC

NAFLD is a slowly progressing condition in humans though patients with advanced NAFLD may develop cirrhosis and hepatocellular carcinoma (HCC, Angulo et al., 2015; Singh et al., 2015). Therefore, one aim of using rodent models is to accelerate the fibrotic and neoplastic process. We used our database of NAFLD models to identify rodent studies reporting the presence of cirrhosis in under 20 weeks (SupTab 2). The majority of these used choline-deficient diets, with or without methionine deficiency. Other models, including diet-induced obesity designs (e.g. 60% kcal high fat diet), were reported to cause advanced fibrosis (>F3), but this was after a duration of 40-50 weeks.

Similarly, HCC develops in these obese rodents models after typically one to two years(Febbraio et al., 2019). Therefore we searched our database to identify models showing HCC at under 30 weeks (SupTab 2). This again included methionine-/choline-deficient models, plus chemically manipulated models, whereby rodents received diethylnitrosamine or streptozocin parenterally.

A further histological feature of specific interest is periportal inflammation. The centrilobular-predominant steatosis and inflammation is the most frequently observed pattern in adults with NAFLD, however the presence of periportal inflammation is associated with more advanced disease in both adults and children(Brunt et al., 2009; Mann et al., 2016). Periportal-predominant steatosis and inflammation is also observed much more frequently in pre-pubescent boys with NAFLD(Schwimmer et al., 2005). Therefore we used the database to identify models that show portal inflammation (SupTab 2) and identified a variety of models, including chemically induced (e.g. monosodium glutamate), dietary (e.g. high fat, high fructose, & high sucrose), and genetic (e.g. galectin-3 knockout). However, portal inflammation was the least reported histological feature (SupFig 2) therefore we suspect that this may be influenced by under-reporting.

### Heterogeneity in nomenclature of dietary models

Use of specialist diets to induce hepatic steatosis, with or without obesity and insulin resistance, has been the mainstay of rodent models of NAFLD(Hebbard and George, 2011). Over 75% of all models in our database include some form of dietary perturbation. However, we observed that there was inconsistent use of terminology when comparing the names of diets and their composition.

For example, 16% (237/1488) models described as ‘high fat diet’ (without specifying additional components), also contained added cholesterol (Figure 3). Overall, 22% (334/1488) of ‘high fat diets’ had some added cholesterol, choline, sucrose, or fructose/glucose.

**Fig. 3:**
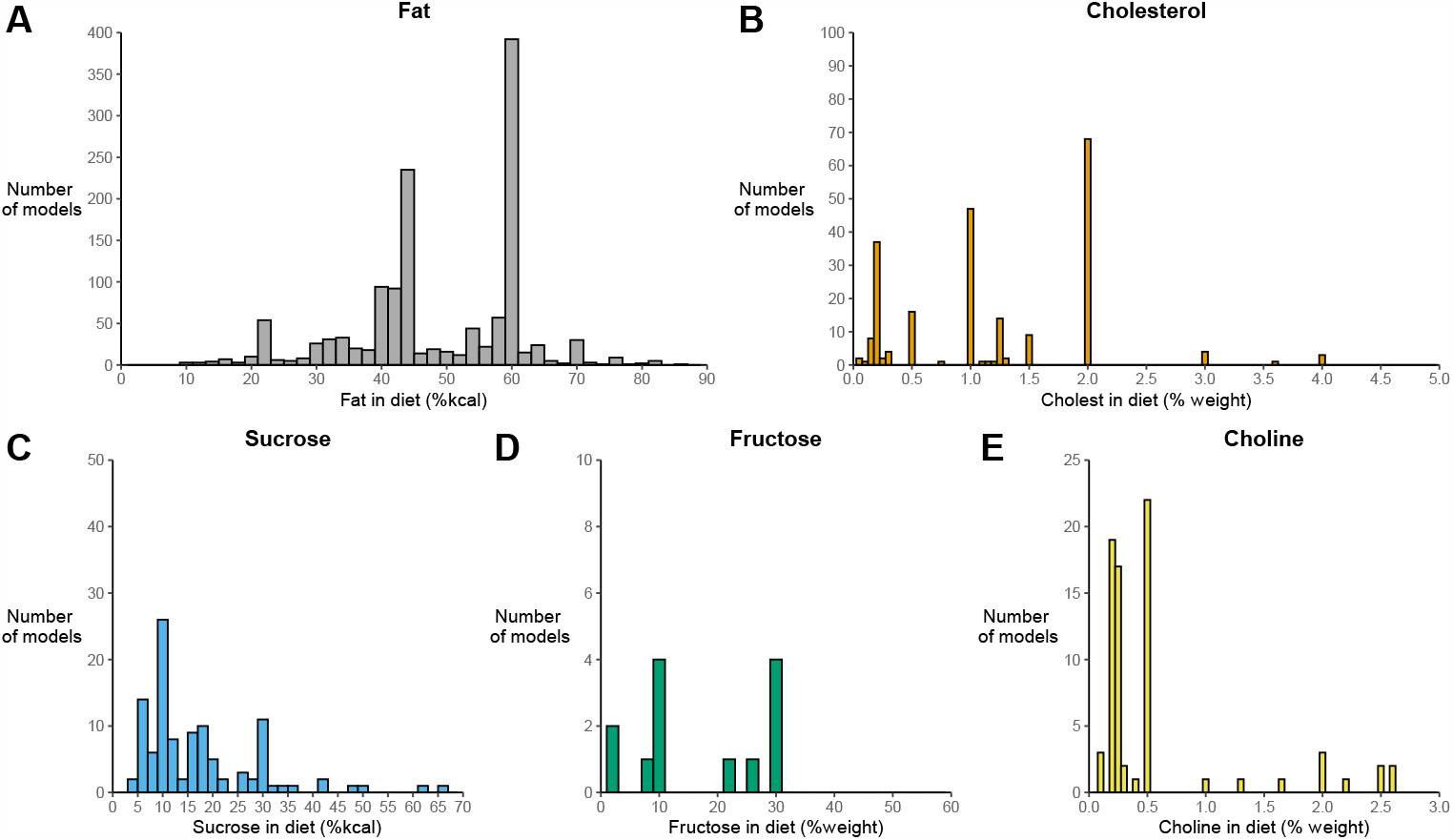
Composition of ‘High fat’ diets. Data from 1488 rodent ‘high fat diet’ models. (A) Proportion of total dietary kcal from fat. (B) Percentage of diet as cholesterol (by weight). (C) Proportion of total dietary kcal from sucrose. (D) Percentage of diet as fructose/glucose (either alone or in combination). (E) Percentage of diet as choline (by weight). The dotted line represents the mean value.

Similar results were observed for 149 ‘Western diet’ models, which most frequently involved 40-45% kcal from fat, plus 30% kcal from sucrose and 0.2% cholesterol. However, there was substantial variability (SupFig 3) with many models described as ‘Western diet’ including added fructose. In addition, 18 models were described as using ‘Western diets’ but provided no detail on diet composition.

### Genetic models provide insight into NAFLD pathogenesis

Understanding the genetics of NAFLD in humans has provided substantial insight into the pathogenesis of fatty liver disease(Romeo et al., 2020). Concurrently, there have been hundreds of genetically modified animal models that display features of NAFLD to varying severity. We used our database to identify 433 human orthologues from genetically modified mice that display exacerbated severity of NAFLD. Using gene set enrichment analysis(Chen et al., 2013; Kuleshov et al., 2016), we identified that these gene targets are most highly expressed in adipose tissue and liver (Figure 4A and SupTable 3). Pathways related to insulin resistance and adipogenesis were enriched (Figure 4B-C), as well genes implicated in type 2 diabetes and related metabolic syndrome traits in humans (Figure 4D). In addition, pathways related to innate immunity (TNF-alpha and IL-6) were strongly enriched in this gene list.

**Fig. 4:**
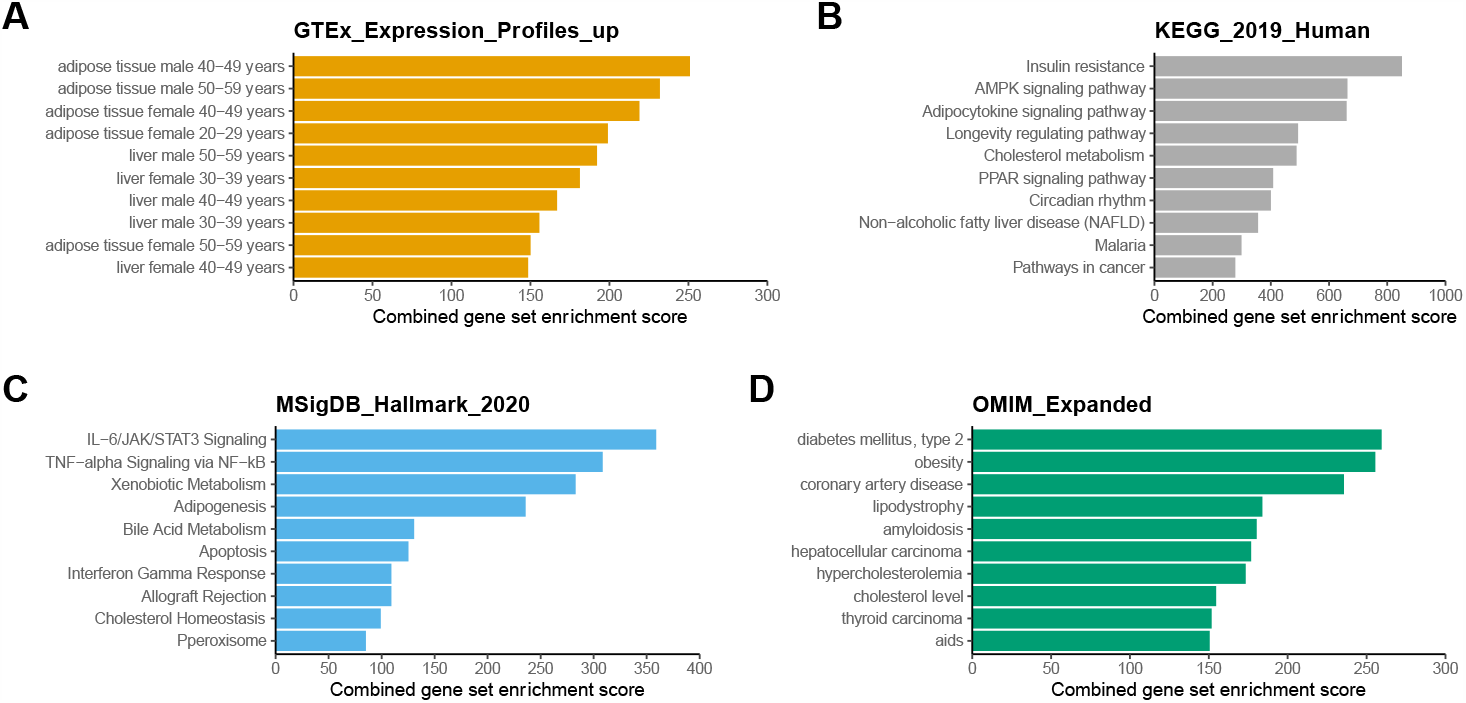
Analysis of genetically modified animals with exacerbated NAFLD. EnrichR gene set enrichment analysis using 433 human orthologues of murine genes from genetically modified animals with increased severity (or susceptibility) to NAFLD. (A) Tissues with high expression of genes (from GTEx). (B) Kyoto Encyclopedia of Genes and Genomes (KEGG) pathway analysis. (C) MSigDB Hallmark Gene Set analysis from Gene Set Enrichment Analysis. (D) Enrichment of genes for human disorders from Online Mendelian Inheritance in Man (OMIM). Related to data in Table S3.

### Risk of bias

Recommendations for reporting of preclinical studies suggest the use of blinding, randomisation, a pre-specified protocol, and a sample size estimate(Percie du Sert et al., 2020). We assessed all 4540 articles for these four risk of bias metrics and combined them into an overall score [0-4]. Over half of all studies had a score of 0 or 1 (SupFig 4). Very few studies reported a power calculation (1.3%), with blinding used in 19% and randomisation in 37%.

## Discussion

Preclinical animal studies of hepatic steatosis have been conducted for nearly 70 years with an aim of helping to elucidate the mechanisms of liver lipid metabolism(Eilert and Dragstedt, 1946), and more recently to find a treatment for NAFLD(Friedman et al., 2018). This has resulted in an increasingly complex array of animal models(Farrell et al., 2019; Nevzorova et al., 2020; Santhekadur et al., 2018). Here, we performed a systematic analysis of animal models of NAFLD, providing a framework for their description, highlighting challenges of reproducibility, and deriving insight into the pathogenesis of NAFLD.

We categorised over 3900 unique models through detailed comparison of animal study design. This large number is due to variation in almost every aspect of design, including age of interventions, dietary composition, sex, and genetic background. Therefore, whilst many studies may initially appear to use the same model (e.g. high fat, high cholesterol in C57BL/6J mice), the majority have only been used by a single study. However, consideration of precise details in study design is relevant in light of recent data that suggests even small differences in age or diet composition may affect treatment response in animal models of NAFLD(Hunter et al., 2020). Moreover, this has implications for understanding the ‘reproducibility crisis’ of preclinical studies(Baker, 2016; Fanelli, 2018; von Herrath et al., 2019). Our results suggest that most models have never been replicated, which adds additional variability in challenging *in vivo* studies with small effect sizes. We found that genetic models were the most frequently replicated and the perturbation is (usually) unambiguously defined (e.g. albumin-Cre liver-specific PTEN knock-out), compared to ‘Western diet’. However, genetically modified animals are subject to compensatory germline expression changes that may affect metabolism through mechanisms that may be difficult to identify.

We observed marked variability in the terminology of dietary interventions. ‘Atherogenic diet’ and ‘Western diet’ have been used in animal studies from the 1950s and 1980s, respectively(Baker et al., 1953; Jaskiewicz et al., 1986), with increasing use since pre-formulated diets became available from major laboratory feed suppliers. Whilst there is some consensus on general composition between suppliers (e.g. atherogenic diets Harlan Teklad TD.88137 and Research Diets D12336), there does not appear to be a formal definition of terminology. From our observations, we suggest the development of consensus recommendations around nomenclature of dietary compositions (e.g. Western diet >30% kcal from fat, >30% kcal from sucrose, no added fructose, <0.5% cholesterol), as this will support consistency and reproducibility.

The integral connection of adipose and liver metabolism is well established and supported by strong human genetic evidence, particular from individuals with lipodystrophy(Lotta et al., 2017; Mann and Savage, 2019). We have replicated this finding through analysis of animal models with exacerbated NAFLD, but in addition identified a role for pathways of innate inflammation (i.e. NF-kB, IL-6, TNF-alpha). Steatohepatitis is a state of sterile inflammation, but to date, all common human genetic variants reproducibly associated with NAFLD at genome-wide significance act via perturbation of lipid metabolism, including variants in or near *PNPLA3, TM6SF2, GCKR, HSD17B13, MBOAT7*, and *MTARC1*(Anstee et al., 2020; Emdin et al., 2020; Mann et al., 2020; Parisinos et al., 2020; Speliotes et al., 2011). The enrichment of inflammatory pathways in our analysis suggests that, across multiple different models, differential activity of the innate immune system can accelerate NASH. There have been candidate studies suggesting that common variants in *MERTK*(Cai et al., 2020) and *IFNL4*(Eslam et al., 2015) are associated with severity of NAFLD in humans, though none of these variants have reached genome wide-significance, unlike for hepatitis C(Ge et al., 2009; Patin et al., 2012). In addition, one recent GWAS using electronic medical records identified a GWAS-significant variant near *IL17RA* associated with NAFLD Activity Score(Namjou et al., 2019). Therefore, it appears that genetic evidence is slowly accumulating for a causal role of innate immune activity in NAFLD.

We have used this dataset to explore whether there is a single animal ‘ideal’ model that reflects the spectrum of human NAFLD, or if different study designs should be used for specific animal phenotypes(Castro and Diehl, 2018; Febbraio et al., 2019; Neff, 2019). Whilst no model demonstrated all features of both histological NAFLD and metabolic syndrome, several studies using a prolonged diet of high fat, high fructose diets had the top overall scores for similarity to human NAFLD(Asgharpour et al., 2016; Dowman et al., 2014), in which fibrosis and HCC development can be expedited by treatment with carbon tetrachloride(Tsuchida et al., 2018). The high fat, high fructose model was identified to have the closest transcriptomic signature to human NAFLD(Teufel et al., 2016). Therefore, this core model design appears to most closely mimic the human phenotype, though few model designs have been precisely replicated, so there is insufficient data to recommend specific dietary composition, genetic background, or timings of interventions.

In this study we have concentrated on disease phenotypes, rather than gene expression profiling to compare models. This has facilitated the inclusion of a large number of studies and allowed focus on end-points that are directly comparable to clinical practice. However, we were limited by the level of detail reported by each individual study, which could have led us to overestimate the number of unique models, for example, where two studies were otherwise identical, if one described use of ‘C57BL/6J’ mice and the other reported ‘C57BL/6’, these would be identified as separate designs. Though the importance of genetic (sub-)strains have been demonstrated by results from the Hybrid Mouse Diversity Panel have illustrated the importance of differences in genetic background in murine NAFLD phenotypes(Chella Krishnan et al., 2018; Hui et al., 2015, 2018; Norheim et al., 2018). Finally, there is a wide range of other variables that can affect the phenotype that we have not included in our analysis, including gut microbiome composition(Lee et al., 2020).

## Conclusion

Due to variation in study design, most rodent models of NAFLD have only been used once. This variation is compounded by a lack of consensus around nomenclature of diets. Genetic models that exacerbate NAFLD are enriched for genes involved in adipogenesis and innate immune pathways. Overall, high fat high fructose diet models show the most phenotypic similarity to human NAFLD.

## Author contributions

YRI, HH, DdGH and AD, performed most of the data extraction and analysis. QC, JD. MF, CH, AL, HKL, LM, CGM, RR, and MU, assisted in article screening, data extraction, and curation. IAR assisted in data interpretation and analysis. JPM conceived the project, guided the research, assisted in data interpretation, and wrote the manuscript. All authors edited and approved the manuscript.

## Experimental procedures

### RESOURCE AVAILABILITY

#### Lead Contact

Further information and requests should be directed to Lead Contact, Jake P. Mann.

#### Materials Availability

This study did not generate new unique reagents.

#### Data and Code Availability

The raw dataset used for analysis, including references to individual studies and R code used for analysis, are available from the Dryad repository: https://doi.org/10.5061/dryad.pnvx0k6kq.

## METHOD DETAILS

### Protocol and search strategy

The systematic review protocol was prospectively registered with SyRF (Systematic Review Facility) in August 2017 and is available from: https://drive.google.com/file/d/0B7Z0eAxKc8ApQ0p4OG5SblRlRTA/view. A partial analysis of this dataset has been reported elsewhere(Hunter et al., 2020).

PubMed via MEDLINE, EMBASE, NCBI Gene Expression Omnibus were searched for published articles of experimental rodent models of fatty liver, NAFLD, or non-alcoholic steatohepatitis (NASH). The search was completed in January 2019.

### Study selection and eligibility criteria

Our inclusion criteria were: primary research articles using mice or rats to model NAFLD (to include hepatic steatosis, NASH, and NASH-fibrosis), evidence of hepatic steatosis in model rodents through either increased hepatic triglyceride content or histological assessment. Studies were excluded if: not modelling NAFLD/NASH; studies in humans or any animal other than mice and rats; reviews, comments, letters, editorials, meta-analyses, ideas; articles not in English (unless there was an available translation).

Abstracts and titles were screened to identify relevant studies using Rayyan(Ouzzani et al., 2016). Potentially relevant studies had their full text extracted and were assessed against inclusion/exclusion criteria independently by two reviewers, with discrepancies settled by discussion with JPM.

### Data collection

The variables extracted were as follows: phenotypic characteristics of animal model used (e.g. sex, dietary composition, rodent age, genetic alterations, background animal strain, chemical agents, genetic manipulations); features of metabolic syndrome (obesity, dyslipidaemia, insulin resistance); presence of lipodystrophy; features of NAFLD (elevated aminotransferases, lobular inflammation, hepatocellular ballooning, portal inflammation, NASH, fibrosis (stage, if stated), hepatocellular carcinoma); age (in weeks) each of these features was reported/described. Where data had been deposited in the Gene Expression Omnibus, the GSE accession number was extracted. Mouse Genome Informatics was used to find human orthologues for murine genes perturbed in genetically modified animals(Eppig et al., 2017).

The presence or absence of phenotypic characteristics was based on the reported description in the included article. Lipodystrophy was defined as the presence of reduced fat mass plus increased insulin resistance and where the animals were described as lipodystrophic. Obesity was defined as a significantly higher body weight (or fat mass) than control animals. Insulin resistance was defined as a significant elevation in any of: fasting or postprandial glucose or insulin, homeostatic model assessment of insulin resistance (HOMA-IR), greater area under the curve on glucose or insulin tolerance test, lower glucose infusion rate during hyperinsulinemic-euglycemic clamp studies. Dyslipidemia was defined by any of: higher triglycerides, lower high-density lipoprotein, higher cholesterol, or higher low-density lipoprotein. Elevated aminotransferases were defined by significantly higher alanine or aspartate aminotransferase compared to control animals. Presence or absence of NASH was assessed dichotomously based on description in the article. Many models were dichotomously described to show a presence/absence of fibrosis without further details, therefore this was collected in addition to fibrosis stage (where available), which was extracted according to Kleiner staging(Kleiner et al., 2005).

Genetically modified animals were included as a separate model where they were shown to exacerbate the liver phenotype of NAFLD. If a study reported a ‘negative’ control animal, a model with NAFLD (‘positive control’), and a genetically modified animal with less severe NAFLD than the ‘positive control’, the genetically modified animal would not be included as a separate model of NAFLD.

Studies frequently included multiple model designs (or treatment arms for interventional studies). Data were extracted for each model or interventional arm separately. Where possible, if data from male and female animals were reported separately, they were included as separate models.

### Comparison of models for similarity

Models were compared for similarity based on study design. Criteria for considering two models to be the same were: identical genetic background, sex of animals studied, age diet initiated ±1 week, %kcal of diet from fat ±2%, %weight of cholesterol in diet ±0.1%, %weight of choline in diet ±0.1%, %kcal of diet from sucrose ±2%, fructose-glucose in diet ±2% kcal (or ±2g/L where dissolved in drinking water), age given chemical/virus/surgery ±1 week, dose of chemical ±10% (chemical-specific), and identical route of administration of chemical. Models could be combined using variables that were not reported (e.g. studies not reporting sex, but otherwise identical models, would be considered the same).

Models were organised into three levels: Category, Main group, and Subgroup. This was based on the similarity of model design after data extraction, rather than from terminology used in the original article. For example, a model using 45% kcal fat plus 20% kcal fructose would be categorised as: ‘Diet only’ -> ‘High fat, high carbohydrate’ -> ‘High fat, high fructose’, even if the authors described the model as ‘high fat diet’. The article’s original description used for each model was retained for comparison of nomenclature with study design.

### QUANTIFICATION AND STATISTICAL ANALYSIS

#### Comparison with human phenotype

A metabolic syndrome score (range 0-3) and a liver histology score (range 0-11) were used to assess the similarity of each model with human NAFLD. For the metabolic syndrome score, models scored 1 for each of: obesity, insulin resistance, and dyslipidemia. For the liver histology, models scored 1 for each of: raised aminotransferases, lobular inflammation, portal inflammation, hepatocellular ballooning, NASH, fibrosis (presence/absence), fibrosis stages 1-4 (one point for each), and HCC.

These two scores were combined to give an overall phenotype score (range 0-14).

#### Risk of bias assessment

Each paper was assessed in the following 4 areas: reporting the presence of a protocol, reporting use of randomisation, reporting use of blinding, and a power calculation for sample size estimation. These were each given a score of 1, and each paper was assigned an overall “risk of bias score”.

#### Statistical analysis

For comparison of phenotype scores by Subgroup, models were filtered for those that had been used by at least one study and then Subgroups were filtered for those that contained at least three models.

Gene lists were analysed using EnrichR package for R(Chen et al., 2013; Kuleshov et al., 2016). Histograms and descriptive statistics were extracted using R 3.6.2 for Mac(Harrer et al., 2019; R Core Team, 2019).

## Supporting information

Raw database

Rcode for analysis

ReadMe for database

## Supplementary items

**SupTab 1**: Genetic backgrounds used in rodent models of NAFLD. Data from 3920 rodent models of NAFLD with the 27 most frequently used genetic backgrounds, divided by model category.

**SupTab 2 [Excel spreadsheet]**: Rodent models of NAFLD with specific characteristics (cirrhosis, HCC, inflammation, lipodystrophy). Models that were used by at least two studies and showed either: cirrhosis (fibrosis stage 4) at <20 weeks, HCC at <30 weeks or periportal inflammation. In addition, lipodystrophic models are listed.

**SupTab 3**: Summary results from gene set enrichment analysis of human orthologues from genetically modified rodents with exacerbated NAFLD. Results from EnrichR analysis of 433 genes.

**Sup Fig. 1**: Age at description of histological features of NAFLD. Histograms illustrating that maximum age that models reported the presence of each histological feature of NAFLD.

**Sup Fig. 2**: Reporting of histological features. Proportion of studies reporting each histological feature in 3657 unique rodent models of NAFLD where histological features were described. ‘Unclear’ refers to conflicting reports of histological features in multiple studies.

**Sup Fig. 3**: Composition of ‘Western’ diets. Data from 149 rodent ‘Western diet’ models. (A) Proportion of total dietary kcal from fat. (B) Percentage of diet as cholesterol (by weight). (C) Proportion of total dietary kcal from sucrose. (D) Percentage of diet as fructose/glucose (either alone or in combination). (E) Percentage of diet as choline (by weight). The dotted line represents the mean value.

**Sup Fig. 4**: Risk of bias of included studies. Studies were assessed for the use of a power calculation, blinding, randomisation, and a pre-specified protocol. Each factor was given a score of 1 to generate an overall risk of bias score of 0-4. (A) Distribution of overall risk of bias scores across 4540 included studies. (B) Proportion of studies meeting each of the bias metrics.

## References

Angulo, P., Kleiner, D.E., Dam-Larsen, S., Adams, L. a., Bjornsson, E.S., Charatcharoenwitthaya, P., Mills, P.R., Keach, J.C., Lafferty, H.D., Stahler, A., et al. (2015). Liver Fibrosis, but No Other Histologic Features, Is Associated With Long-term Outcomes of Patients With Nonalcoholic Fatty Liver Disease. Gastroenterology 149, 389–397.e10.

Anstee, Q.M., Darlay, R., Cockell, S., Meroni, M., Govaere, O., Tiniakos, D., Burt, A.D., Bedossa, P., Palmer, J., Liu, Y.-L., et al. (2020). Genome-wide association study of non-alcoholic fatty liver and steatohepatitis in a histologically-characterised cohort. J. Hepatol.

Asgharpour, A., Cazanave, S.C., Pacana, T., Seneshaw, M., Vincent, R., Banini, B.A., Kumar, D.P., Daita, K., Min, H.-K., Mirshahi, F., et al. (2016). A diet-induced animal model of non-alcoholic fatty liver disease and hepatocellular cancer. J. Hepatol. 65, 579–588.

Baker, M. (2016). Reproducibility crisis. Nature 533, 353–366.

Baker, S.P., Ogden, E., and Riddle, J.W. (1953). Serological detection of abnormal concentrations of serum lipoproteins in dogs on cholesterol-thiouracil atherogenic diets. Proc. Soc. Exp. Biol. Med. 82, 119–122.

Brunt, E.M., Kleiner, D.E., Wilson, L. a., Unalp, A., Behling, C.E., Lavine, J.E., Neuschwander-Tetri, B.A., and NASH Clinical Research Network (2009). Portal chronic inflammation in nonalcoholic fatty liver disease (NAFLD): a histologic marker of advanced NAFLD-clinicopathologic correlation from the Nonalcoholic Steatohepatitis Clinical Research Network. Hepatology 49, 809–820.

Brunt, E.M., Kleiner, D.E., Carpenter, D.H., Rinella, M., Harrison, S.A., Loomba, R., Younossi, Z., Neuschwander-Tetri, B.A., Sanyal, A.J., and American Association for the Study of Liver Diseases NASH Task Force (2020). Nonalcoholic fatty liver disease: Reporting histologic findings in clinical practice. Hepatology.

Cai, B., Dongiovanni, P., Corey, K.E., Wang, X., Shmarakov, I.O., Zheng, Z., Kasikara, C., Davra, V., Meroni, M., Chung, R.T., et al. (2020). Macrophage MerTK Promotes Liver Fibrosis in Nonalcoholic Steatohepatitis. Cell Metab. 31, 406–421.e7.

Castro, R.E., and Diehl, A.M. (2018). Towards a definite mouse model of NAFLD. Journal of Hepatology 69, 272–274.

Chella Krishnan, K., Kurt, Z., Barrere-Cain, R., Sabir, S., Das, A., Floyd, R., Vergnes, L., Zhao, Y., Che, N., Charugundla, S., et al. (2018). Integration of Multi-omics Data from Mouse Diversity Panel Highlights Mitochondrial Dysfunction in Non-alcoholic Fatty Liver Disease. Cell Systems 6, 103–115.e7.

Chen, E.Y., Tan, C.M., Kou, Y., Duan, Q., Wang, Z., Meirelles, G.V., Clark, N.R., and Ma’ayan, A. (2013). Enrichr: interactive and collaborative HTML5 gene list enrichment analysis tool. BMC Bioinformatics 14, 128.

Diehl, A.M., and Day, C. (2017). Cause, Pathogenesis, and Treatment of Nonalcoholic Steatohepatitis. N. Engl. J. Med. 377, 2063–2072.

Dongiovanni, P., Stender, S., Pietrelli, A., Mancina, R.M., Cespiati, A., Petta, S., Pelusi, S., Pingitore, P., Badiali, S., Maggioni, M., et al. (2018). Causal relationship of hepatic fat with liver damage and insulin resistance in nonalcoholic fatty liver. J. Intern. Med. 283, 356–370.

Dowman, J.K., Hopkins, L.J., Reynolds, G.M., Nikolaou, N., Armstrong, M.J., Shaw, J.C., Houlihan, D.D., Lalor, P.F., Tomlinson, J.W., Hübscher, S.G., et al. (2014). Development of hepatocellular carcinoma in a murine model of nonalcoholic steatohepatitis induced by use of a high-fat/fructose diet and sedentary lifestyle. Am. J. Pathol. 184, 1550–1561.

Eilert, M.L., and Dragstedt, L.R. (1946). Lipotropic action of lipocaic; a study of the effect of oral and parenteral lipocaic and oral inositol on the dietary fatty liver of the white rat. Am. J. Physiol. 147, 346–351.

Emdin, C.A., Haas, M.E., Khera, A.V., Aragam, K., Chaffin, M., Klarin, D., Hindy, G., Jiang, L., Wei, W.-Q., Feng, Q., et al. (2020). A missense variant in Mitochondrial Amidoxime Reducing Component 1 gene and protection against liver disease. PLoS Genet. 16, e1008629.

Eppig, J.T., Smith, C.L., Blake, J.A., Ringwald, M., Kadin, J.A., Richardson, J.E., and Bult, C.J. (2017). Mouse Genome Informatics (MGI): Resources for Mining Mouse Genetic, Genomic, and Biological Data in Support of Primary and Translational Research. In Systems Genetics: Methods and Protocols, K. Schughart, and R.W. Williams, eds. (New York, NY: Springer New York), pp. 47–73.

Eslam, M., Hashem, A.M., Leung, R., Romero-Gomez, M., Berg, T., Dore, G.J., Chan, H.L.K., Irving, W.L., Sheridan, D., Abate, M.L., et al. (2015). Interferon-λ rs12979860 genotype and liver fibrosis in viral and non-viral chronic liver disease. Nat. Commun. 6, 6422.

Eslam, M., Sanyal, A.J., George, J., and International Consensus Panel (2020). MAFLD: A Consensus-Driven Proposed Nomenclature for Metabolic Associated Fatty Liver Disease. Gastroenterology 158, 1999–2014.e1.

Fanelli, D. (2018). Opinion: Is science really facing a reproducibility crisis, and do we need it to? Proceedings of the National Academy of Sciences 115, 2628–2631.

Farrell, G., Schattenberg, J.M., Leclercq, I., Yeh, M.M., Goldin, R., Teoh, N., and Schuppan, D. (2019). Mouse Models of Nonalcoholic Steatohepatitis: Toward Optimization of Their Relevance to Human Nonalcoholic Steatohepatitis. Hepatology 69, 2241–2257.

Febbraio, M.A., Reibe, S., Shalapour, S., Ooi, G.J., Watt, M.J., and Karin, M. (2019). Preclinical Models for Studying NASH-Driven HCC: How Useful Are They? Cell Metab. 29, 18–26.

Friedman, S.L., Neuschwander-Tetri, B.A., Rinella, M., and Sanyal, A.J. (2018). Mechanisms of NAFLD development and therapeutic strategies. Nat. Med. 24, 908– 922.

Ge, D., Fellay, J., Thompson, A.J., Simon, J.S., Shianna, K.V., Urban, T.J., Heinzen, E.L., Qiu, P., Bertelsen, A.H., Muir, A.J., et al. (2009). Genetic variation in IL28B predicts hepatitis C treatment-induced viral clearance. Nature 461, 399–401.

Harrer, M., Cuijpers, P., Furukawa, T.A., and Ebert, D.D. (2019). Doing Meta-Analysis in R: A Hands-on Guide. PROTECT Lab Erlangen.

Hebbard, L., and George, J. (2011). Animal models of nonalcoholic fatty liver disease. Nat. Rev. Gastroenterol. Hepatol. 8, 35–44.

von Herrath, M., Pagni, P.P., Grove, K., Christoffersson, G., Tang-Christensen, M., Karlsen, A.E., and Petersen, J.S. (2019). Case Reports of Pre-clinical Replication Studies in Metabolism and Diabetes. Cell Metab. 29, 795–802.

Hui, S.T., Parks, B.W., Org, E., Norheim, F., Che, N., Pan, C., Castellani, L.W., Charugundla, S., Dirks, D.L., Psychogios, N., et al. (2015). The genetic architecture of NAFLD among inbred strains of mice. Elife 4, e05607.

Hui, S.T., Kurt, Z., Tuominen, I., Norheim, F.C Davis, R., Pan, C., Dirks, D.L., Magyar, C.E., French, S.W., Chella Krishnan, K., et al. (2018). The Genetic Architecture of Diet-Induced Hepatic Fibrosis in Mice. Hepatology 68, 2182–2196.

Hunter, H., de Gracia Hahn, D., Duret, A., Im, Y.R., Cheah, Q., Dong, J., Fairey, M., Hjalmarsson, C., Li, A., Lim, H.K., et al. (2020). Weight loss, insulin resistance, and study design confound results in a meta-analysis of animal models of fatty liver. Elife 9.

Jaskiewicz, K., Rossouw, J.E., Kritchevsky, D., van Rensburg, S.J., Fincham, J.E., and Woodroof, C.W. (1986). The influence of diet and dimethylhydrazine on the small and large intestine of vervet monkeys. Br. J. Exp. Pathol. 67, 361–369.

Kleiner, D.E., Brunt, E.M., Van Natta, M., Behling, C., Contos, M.J., Cummings, O.W., Ferrell, L.D., Liu, Y.-C., Torbenson, M.S., Unalp-Arida, A., et al. (2005). Design and validation of a histological scoring system for nonalcoholic fatty liver disease. Hepatology 41, 1313–1321.

Kuleshov, M.V., Jones, M.R., Rouillard, A.D., Fernandez, N.F., Duan, Q., Wang, Z., Koplev, S., Jenkins, S.L., Jagodnik, K.M., Lachmann, A., et al. (2016). Enrichr: a comprehensive gene set enrichment analysis web server 2016 update. Nucleic Acids Res. 44, W90–W97.

Lee, G., You, H.J., Bajaj, J.S., Joo, S.K., Yu, J., Park, S., Kang, H., Park, J.H., Kim, J.H., Lee, D.H., et al. (2020). Distinct signatures of gut microbiome and metabolites associated with significant fibrosis in non-obese NAFLD. Nat. Commun. 11, 4982.

Liu, Z., Zhang, Y., Graham, S., Wang, X., Cai, D., Huang, M., Pique-Regi, R., Dong, X.C., Chen, Y.E., Willer, C., et al. (2020). Causal relationships between NAFLD, T2D and obesity have implications for disease subphenotyping. J. Hepatol.

Lotta, L.A., Gulati, P., Day, F.R., Payne, F., Ongen, H., van de Bunt, M., Gaulton, K.J., Eicher, J.D., Sharp, S.J., Luan J. ‘an, et al. (2017). Integrative genomic analysis implicates limited peripheral adipose storage capacity in the pathogenesis of human insulin resistance. Nat. Genet. 49, 17–26.

Luukkonen, P.K., Zhou, Y., Sädevirta, S., Leivonen, M., Arola, J., Orešič, M., Hyötyläinen, T., and Yki-Järvinen, H. (2016). Hepatic ceramides dissociate steatosis and insulin resistance in patients with non-alcoholic fatty liver disease. J. Hepatol. 64, 1167–1175.

Mann, J.P., and Savage, D.B. (2019). What lipodystrophies teach us about the metabolic syndrome. J. Clin. Invest. 130, 4009–4021.

Mann, J.P., Vito, R.D., Mosca, A., Alisi, A., Armstrong, M.J., Raponi, M., Baumann, U., and Nobili, V. (2016). Portal inflammation is independently associated with fibrosis and metabolic syndrome in pediatric nonalcoholic fatty liver disease. Hepatology 63, 745– 753.

Mann, J.P., Pietzner, M., Wittemans, L.B., De Lucia Rolfe, E., Nicola, D., Imamura, F., Forouhi, N.G., Fauman, E., Allison, M.E., Jules, L., et al. (2020). Insights into genetic variants associated with NASH-fibrosis from metabolite profiling. Hum Mol Genet doi: 10.1093/hmg/ddaa162.

Namjou, B., Lingren, T., Huang, Y., Parameswaran, S., Cobb, B.L., Stanaway, I.B., Connolly, J.J., Mentch, F.D., Benoit, B., Niu, X., et al. (2019). GWAS and enrichment analyses of non-alcoholic fatty liver disease identify new trait-associated genes and pathways across eMERGE Network. BMC Med. 17, 135.

Neff, E.P. (2019). Farewell, FATZO: a NASH mouse update. Lab Animal 48, 151.

Nevzorova, Y.A., Boyer-Diaz, Z., Cubero, F.J., and Gracia-Sancho, J. (2020). Animal models for liver disease - A practical approach for translational research. J. Hepatol. 73, 423–440.

Norheim, F., Bjellaas, T., Hui, S.T., Chella Krishnan, K., Lee, J., Gupta, S., Pan, C., Hasin-Brumshtein, Y., Parks, B.W., Li, D.Y., et al. (2018). Genetic, dietary, and sex-specific regulation of hepatic ceramides and the relationship between hepatic ceramides and IR. J. Lipid Res. 59, 1164–1174.

Ouzzani, M., Hammady, H., Fedorowicz, Z., and Elmagarmid, A. (2016). Rayyan-a web and mobile app for systematic reviews. Syst. Rev. 5, 210.

Parisinos, C.A., Wilman, H.R., Thomas, E.L., Kelly, M., Nicholls, R.C., McGonigle, J., Neubauer, S., Hingorani, A.D., Patel, R.S., Hemingway, H., et al. (2020). Genome-wide and Mendelian randomisation studies of liver MRI yield insights into the pathogenesis of steatohepatitis. J. Hepatol. 73, 241–251.

Patin, E., Kutalik, Z., Guergnon, J., Bibert, S., Nalpas, B., Jouanguy, E., Munteanu, M., Bousquet, L., Argiro, L., Halfon, P., et al. (2012). Genome-wide association study identifies variants associated with progression of liver fibrosis from HCV infection. Gastroenterology 143, 1244–1252.e12.

Percie du Sert, N., Hurst, V., Ahluwalia, A., Alam, S., Avey, M.T., Baker, M., Browne, W.J., Clark, A., Cuthill, I.C., Dirnagl, U., et al. (2020). The ARRIVE guidelines 2.0: Updated guidelines for reporting animal research. PLoS Biol. 18, e3000410.

R Core Team (2019). A language and environment for statistical computing. Vienna, Austria: R Foundation for Statistical Computing.

Romeo, S., Sanyal, A., and Valenti, L. (2020). Leveraging Human Genetics to Identify Potential New Treatments for Fatty Liver Disease. Cell Metab. 31, 35–45.

Samuel, V.T., and Shulman, G.I. (2018). Nonalcoholic Fatty Liver Disease as a Nexus of Metabolic and Hepatic Diseases. Cell Metab. 27, 22–41.

Santhekadur, P.K., Kumar, D.P., and Sanyal, A.J. (2018). Preclinical models of non-alcoholic fatty liver disease. J. Hepatol. 68, 230–237.

Schwimmer, J.B., Behling, C., Newbury, R., Deutsch, R., Nievergelt, C., Schork, N.J., and Lavine, J.E. (2005). Histopathology of pediatric nonalcoholic fatty liver disease. Hepatology 42, 641–649.

Singh, S., Allen, A.M., Wang, Z., Prokop, L.J., Murad, M.H., and Loomba, R. (2015). Fibrosis Progression in Nonalcoholic Fatty Liver vs Nonalcoholic Steatohepatitis: A Systematic Review and Meta-analysis of Paired-Biopsy Studies. Clinical Gastroenterology and Hepatology 13, 643–654.

Speliotes, E.K., Yerges-armstrong, L.M., Wu, J., Hernaez, R., Lauren, J., Palmer, C.D., Gudnason, V., Eiriksdottir, G., Garcia, M.E., Launer, L.J., et al. (2011). Genome-Wide Association Analysis Identifies Variants Associated with Nonalcoholic Fatty Liver Disease That Have Distinct Effects on Metabolic Traits. PLoS Genet. 7, e1001324.

Teufel, A., Itzel, T., Erhart, W., Brosch, M., Wang, X.Y., Kim, Y.O., von Schönfels, W., Herrmann, A., Brückner, S., Stickel, F., et al. (2016). Comparison of Gene Expression Patterns Between Mouse Models of Nonalcoholic Fatty Liver Disease and Liver Tissues From Patients. Gastroenterology 151, 513–525.e0.

Tsuchida, T., Lee, Y.A., Fujiwara, N., Ybanez, M., Allen, B., Martins, S., Fiel, M.I., Goossens, N., Chou, H.-I., Hoshida, Y., et al. (2018). A simple diet- and chemical-induced murine NASH model with rapid progression of steatohepatitis, fibrosis and liver cancer. J. Hepatol. 69, 385–395.

