## Supplementary data for "A systematic analysis of rodent models implicates adipogenesis and innate immunity in pathogenesis of fatty liver disease"

### Supplementary items

| Genetic background | Overall | Dietary | Dietary + other | Genetic | Genetic + chemical | Genetic + dietary | Genetic + other | Chemical | Offspring | Other |
|---|---|---|---|---|---|---|---|---|---|---|
| 129-C57BL/6 | 103 (2.6%) | 27 (1.4%) | 1 (1.5%) | 40 (8.4%) | 1 (1.5%) | 33 (4%) | | 1 (0.3%) | | |
| 129Sv | 56 (1.4%) | 25 (1.3%) | | 11 (2.3%) | 2 (3.1%) | 12 (1.5%) | 2 (7.7%) | 4 (1%) | | |
| A/J | 16 (0.4%) | 14 (0.7%) | | | | 1 (0.1%) | | 1 (0.3%) | | |
| BALB/c | 51 (1.3%) | 26 (1.3%) | 2 (2.9%) | 5 (1%) | | 6 (0.7%) | | 11 (2.8%) | | 1 (2%) |
| BALB/c-C57BL/6 | 5 (0.1%) | 2 (0.1%) | | 2 (0.4%) | | 1 (0.1%) | | | | |
| C3H | 37 (0.9%) | 30 (1.6%) | | 6 (1.3%) | | | | 1 (0.3%) | | |
| C57BL/6-C3H | 4 (0.1%) | 2 (0.1%) | | 1 (0.2%) | | 1 (0.1%) | | | | |
| C57BL/6-Other | 94 (2.4%) | 42 (2.2%) | | 27 (5.7%) | 6 (9.2%) | 15 (1.8%) | | 4 (1%) | | |
| C57BL/6? | 693 (17.7%) | 298 (15.5%) | 11 (16.2%) | 102 (21.4%) | 15 (23.1%) | 189 (22.9%) | 2 (7.7%) | 59 (15.1%) | 8 (8.8%) | 9 (18.4%) |
| C57BL/6J | 1055 (26.9%) | 457 (23.7%) | 22 (32.4%) | 133 (27.9%) | 18 (27.7%) | 306 (37%) | 12 (46.2%) | 81 (20.7%) | 17 (18.7%) | 9 (18.4%) |
| C57BL/6N | 87 (2.2%) | 55 (2.9%) | | 8 (1.7%) | | 10 (1.2%) | | 14 (3.6%) | | |
| CD-1 | 18 (0.5%) | 8 (0.4%) | | 2 (0.4%) | | 2 (0.2%) | | 5 (1.3%) | 1 (1.1%) | |
| Fischer 344 | 42 (1.1%) | 27 (1.4%) | 2 (2.9%) | 1 (0.2%) | | | | 10 (2.6%) | | 2 (4.1%) |
| FVB | 35 (0.9%) | 15 (0.8%) | 1 (1.5%) | 10 (2.1%) | 1 (1.5%) | 5 (0.6%) | | 3 (0.8%) | | |
| FVB-C57BL/6 | 12 (0.3%) | 4 (0.2%) | | 4 (0.8%) | | 4 (0.5%) | | | | |
| Holtzman | 13 (0.3%) | 12 (0.6%) | | | | | | | 1 (1.1%) | |
| ICR | 53 (1.4%) | 27 (1.4%) | | | | 8 (1%) | | 13 (3.3%) | 3 (3.3%) | 2 (4.1%) |
| KK-Ay | 22 (0.6%) | 3 (0.2%) | | 2 (0.4%) | 2 (3.1%) | 15 (1.8%) | | | | |
| Lewis | 16 (0.4%) | 16 (0.8%) | | | | | | | | |
| Long Evans | 24 (0.6%) | 10 (0.5%) | | 2 (0.4%) | 2 (3.1%) | 9 (1.1%) | | 1 (0.3%) | | |
| NOD.B10 | 18 (0.5%) | 5 (0.3%) | | 4 (0.8%) | | 9 (1.1%) | | | | |
| Not stated | 266 (6.8%) | 61 (3.2%) | 1 (1.5%) | 63 (13.2%) | 10 (15.4%) | 105 (12.7%) | 7 (26.9%) | 15 (3.8%) | 2 (2.2%) | 2 (4.1%) |
| Other | 209 (5.3%) | 85 (4.4%) | 3 (4.4%) | 39 (8.2%) | 3 (4.6%) | 49 (5.9%) | 1 (3.8%) | 20 (5.1%) | 3 (3.3%) | 6 (12.2%) |
| SHR | 18 (0.5%) | 2 (0.1%) | | 4 (0.8%) | | 12 (1.5%) | | | | |

| | | | | | | | | | | |
|---|---|---|---|---|---|---|---|---|---|---|
| Sprague Dawley | 488 (12.4%) | 355 (18.4%) | 15 (22.1%) | 3 (0.6%) | | 8 (1%) | | 80 (20.5%) | 18 (19.8%) | 9 (18.4%) |
| Swiss | 21 (0.5%) | 13 (0.7%) | 2 (2.9%) | | | 1 (0.1%) | | 4 (1%) | 1 (1.1%) | |
| Wistar | 425 (10.8%) | 304 (15.8%) | 7 (10.3%) | 2 (0.4%) | | 5 (0.6%) | | 62 (15.9%) | 36 (39.6%) | 9 (18.4%) |
| Zucker | 39 (1%) | 2 (0.1%) | 1 (1.5%) | 6 (1.3%) | 5 (7.7%) | 20 (2.4%) | 2 (7.7%) | 2 (0.5%) | 1 (1.1%) | |

***SupTab 1**: Genetic backgrounds used in rodent models of NAFLD. Data from 3920 rodent models of NAFLD with the 27 most frequently used genetic backgrounds, divided by model category.*

***SupTab 2 [Excel spreadsheet]:** Rodent models of NAFLD with specific characteristics (cirrhosis, HCC, portal inflammation, lipodystrophy). Models that were used by at least two studies and showed either: cirrhosis (fibrosis stage 4) at <20 weeks, HCC at <30 weeks or periportal inflammation. In addition, lipodystrophic models are listed.*

***SupTab 3 [Excel spreadsheet]:** Summary results from gene set enrichment analysis of human orthologues from genetically modified rodents with exacerbated NAFLD. Results from EnrichR analysis of 433 genes.*

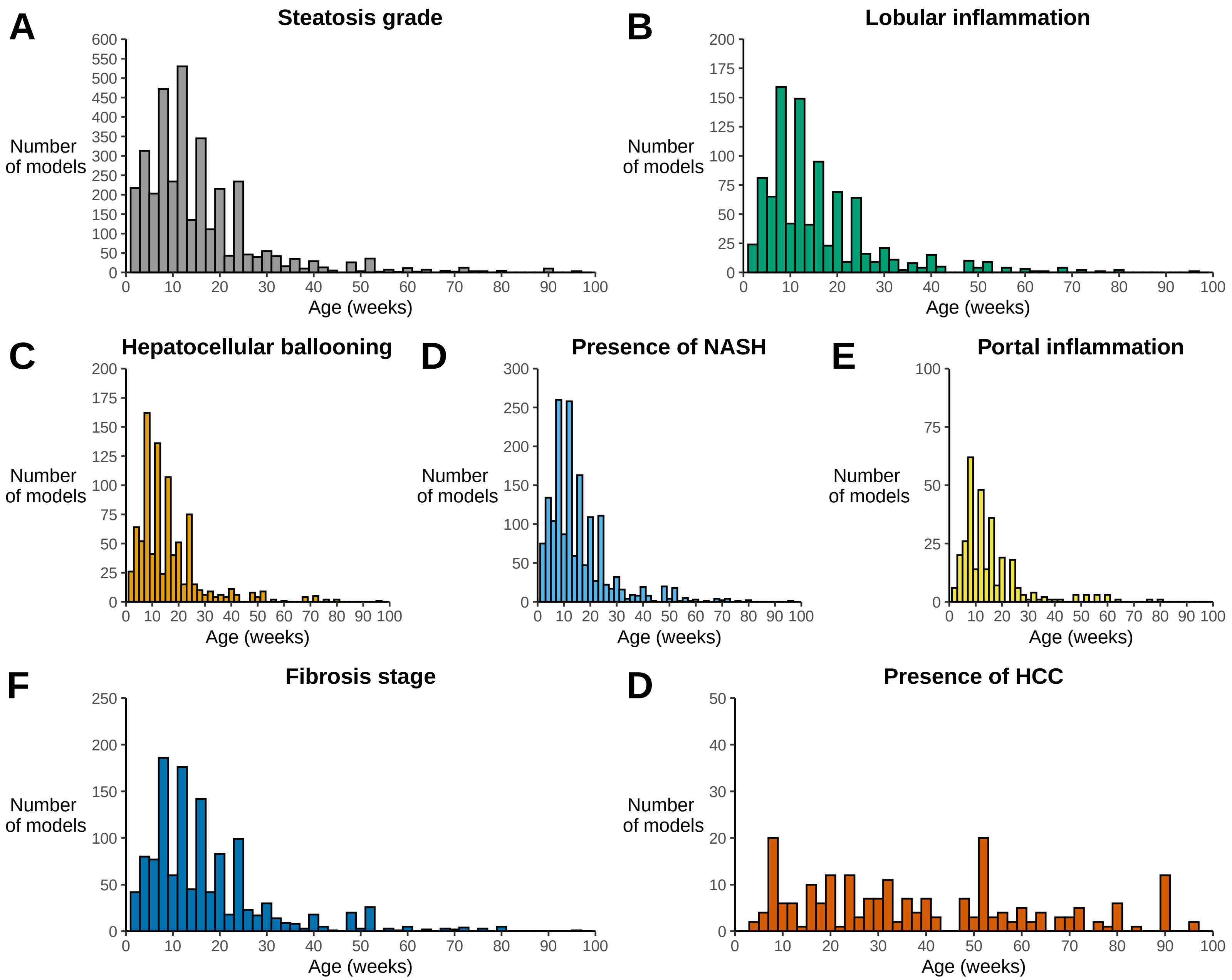


***SupFig. 1:*** *Age at description of histological features of NAFLD. Histograms illustrating that maximum age that models reported the presence of each histological feature of NAFLD.*

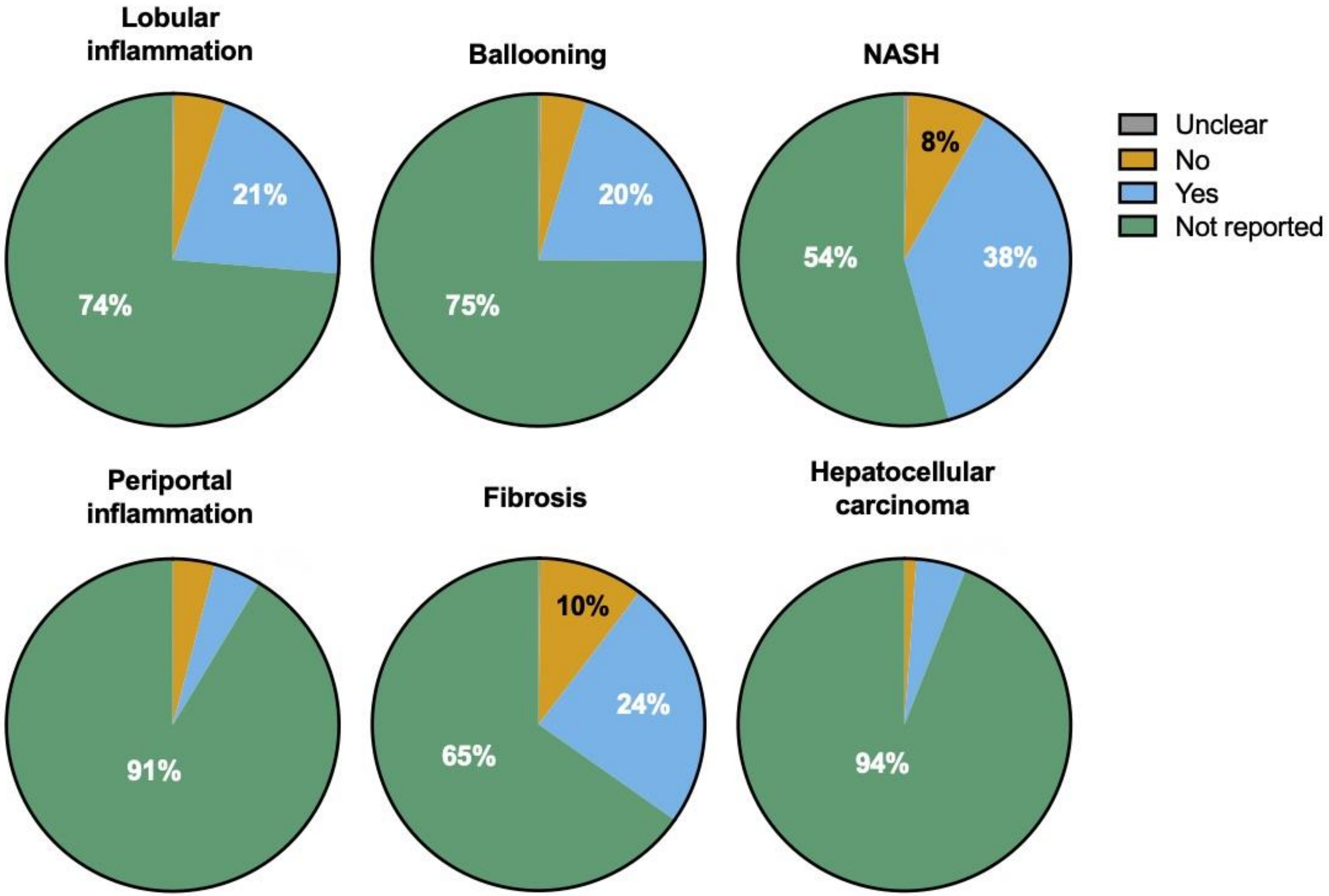


***SupFig. 2:*** *Reporting of histological features. Proportion of studies reporting each histological feature in 3657 unique rodent models of NAFLD where histological features were described. 'Unclear' refers to conflicting reports of histological features in multiple studies.*

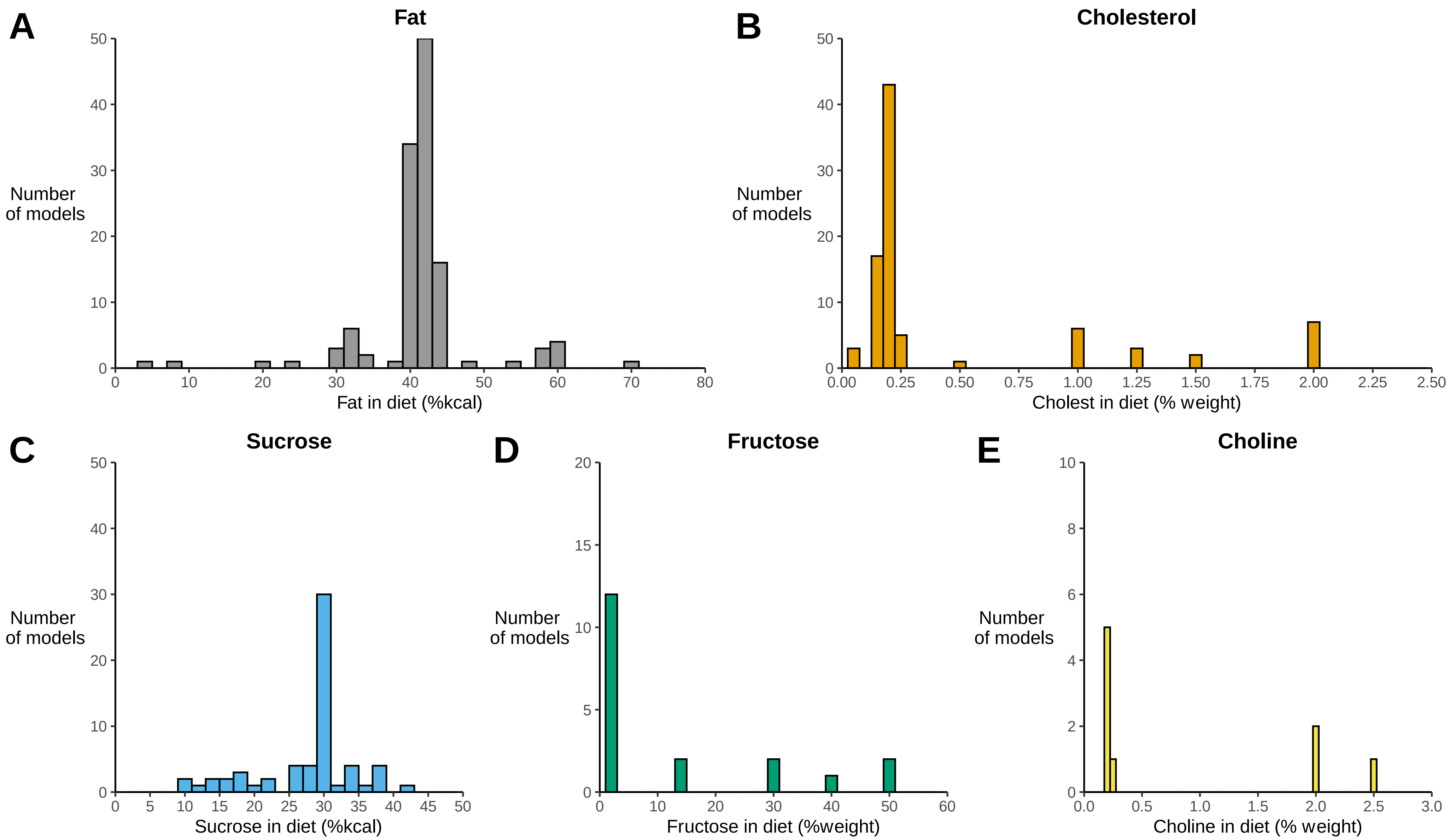


***SupFig. 3**: Composition of 'Western' diets. Data from 149 rodent 'Western diet' models. (A) Proportion of total dietary kcal from fat. (B) Percentage of diet as cholesterol (by weight). (C) Proportion of total dietary kcal from sucrose. (D) Percentage of diet as fructose/glucose (either alone or in combination). (E) Percentage of diet as choline (by weight). The dotted line represents the mean value.*

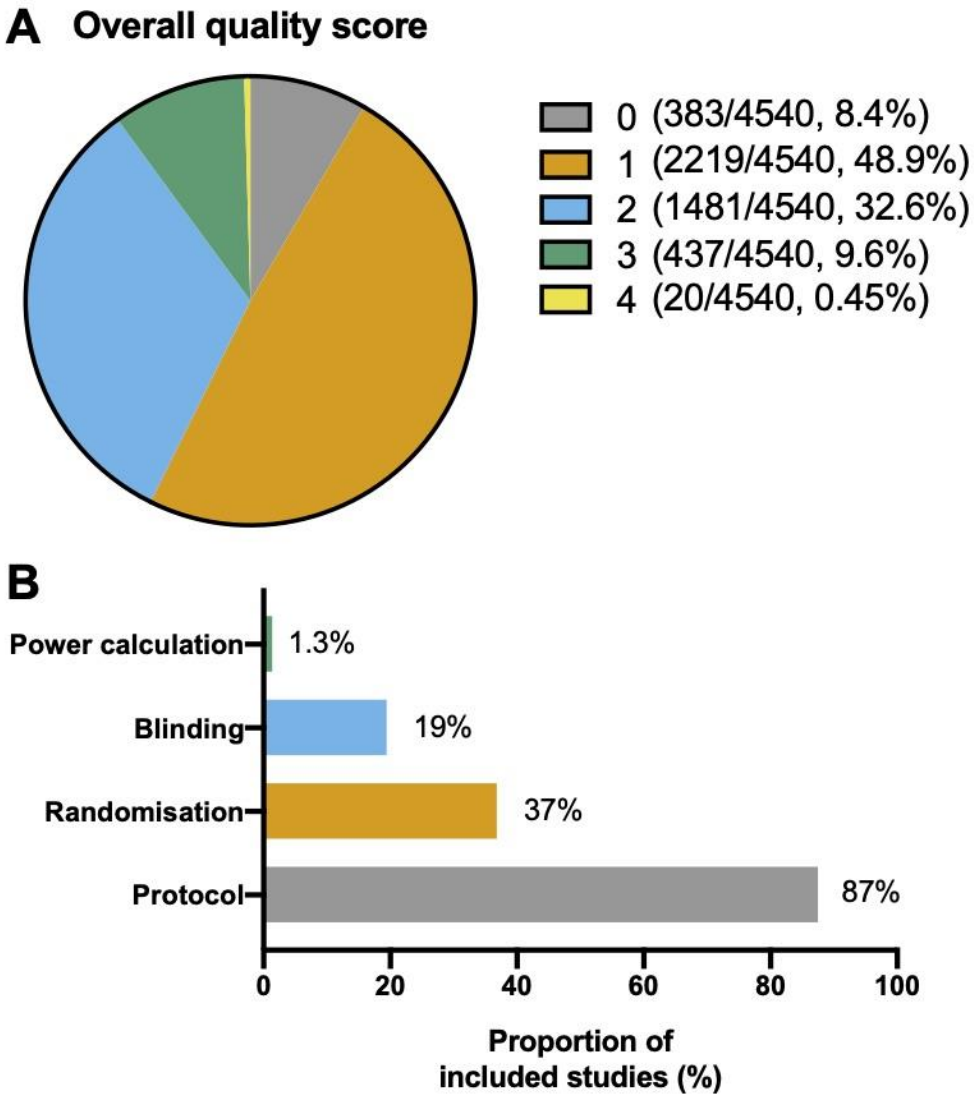


***SupFig. 4****: Risk of bias of included studies. Studies were assessed for the use of a power calculation, blinding, randomisation, and a pre-specified protocol. Each factor was given a score of 1 to generate an overall risk of bias score of 0-4. (A) Distribution of overall risk of bias scores across 4540 included studies. (B) Proportion of studies meeting each of the bias metrics.*
