## Supplementary material for "A systematic analysis of rodent models implicates adipogenesis and innate immunity in pathogenesis of fatty liver disease": ReadMe for database

ReadMe file for NAFLD Animal Models Database v1

Contact:
Jake Mann, Institute of Metabolic Science, Cambridge, UK


Data used in a manuscript currently under review.
Derived from a systematic review of rodent models of NAFLD. Data representes the raw data and references to all included studies.
Search completed January 2019. Data updated November 2020.

The dataset is two tabs in a .xlsx file. The first tab "Papers & refs" gives the references for each article included in the review. The second tab "Models" provides full details of each model included in the review.

The data is free to use with no restrictions.

Full description of the analysis, methods, and processing of the data will be available in the manuscript once published.
The code used for analysis (with R 4.0.2) accompanies this dataset and ReadMe. Note that for analysis Tab1 ('Papers & refs') needs to separated and renamed as "AnMod_papers_Nov20_v2.xlsx"; and Tab2 ('Models') needs to be separated and renamed as "AnMod_models_Nov20_v2.xlsx".


Variable list:
Tab1 "Papers & refs"		
ID	-	Unique study ID. This relates to column BP in the "Models" tab and is used as a the 'Reference ID number' 
Study	-	Surname and Year of the article 
Year 	-	Year article was published 
Title	-	Title of the article included 
Include	-	Include/Exclude against the review's criteria. [Note that excluded studies have not been included in this dataset.] 
Models 	-	The model(s) used by each study. If there are multiple models this will result in multiple rows per study. 
Background	-	Genetic background used in each model. 
Paper_type	-	The paper type, categorised as: Phase 2/3 (i.e. an interventional study using an agent also used in phase 2/3 trials in humans), Mechanistic, Model description, Interventional (other) (i.e. an interventional study not covered by Phase 2/3), or Other 
Other	-	The agent used as part of 'Interventional (other)' studies 
Gene1	-	Key gene1 reported in the abstract of the paper 
Gene2	-	Key gene2 reported in the abstract of the paper 
Gene3	-	Key gene3 reported in the abstract of the paper 
Other mechanistic factor	-	Any other mechanistic factor reported in the abstract of the paper 
Tx_omic	-	GEO dataset GSE code (to be used at: https://www.ncbi.nlm.nih.gov/gds) for transcriptomic data 
Protocol	-	Presence of a protocol 
Randomisation	-	Use of randomisation 
Blinding	-	Use of blinding 
Power_calc	-	Description of a power calculation or sample size estimation 
Qual_score	-	Score (0-4) using the previous four risk of bias items 


		
Tab2 "Models"		 
Category	-	Overall category of the model 
Main_group	-	Main group of the model 
Gene	-	Murine gene involved in genetically modified animals 
HUGO_gene	-	Human ortholog in HUGO format for genetic models 
Subgroup	-	Subgroup for each model 
Detail	-	Description/nomenclature of the model as used by the article (i.e. this has not been edited/curated by us) 
Background	-	Full/long genetic background of the model 
Background_simple	-	Simplified/grouped genetic background 
Sex	-	Sex of animals used 
Diet_detail	-	Age (weeks of life) of when the diet was initiated (blank if not stated) 
Diet_num	-	Age (weeks of life) of when the diet was initiated (blank if not stated) [formatted as number for analysis]
Fat_detail	-	Proportion of fat in diet (%kcal) 
Fat_num	-	Proportion of fat in diet (%kcal) [formatted as number for analysis]
Cholest_per	-	Proportion of cholesterol in diet (%weight) 
Cholest_num	-	Proportion of cholesterol in diet (%weight) [formatted as number for analysis]
Choline_num	-	Proportion of choline in diet (%weight) [formatted as number for analysis]
Sucrose_kcal	-	Proportion of sucrose in diet (%kcal) 
Sucrose_num	-	Proportion of sucrose in diet (%kcal) [formatted as number for analysis]
Fruct_wt	-	Proportion of fructose in diet (%weight) 
Fruct_water_solid	-	Whether fructose was provided as solid or liquid/in water 
Fruct_water_num	-	Proportion of fructose in diet (%weight) where included in water or as a liquid 
Fruct_num	-	Proportion of fructose in diet (%weight) [formatted as number for analysis]
Fruct_solid_num	-	Proportion of fructose in diet (%weight) where included as solid diet 
Chem_age	-	Age that a chemical or surgical intervention was made (in weeks), also to include age that viral agents were given in genetically modified animals 
Chem_age_num	-	Age that a chemical or surgical intervention was made (in weeks), also to include age that viral agents were given in genetically modified animals [formatted as number for analysis]
Chem_dose	-	Dose of chemical agents 
Chem_route_full	-	Full/long route of chemical dose administration 
Chem_route_simple	-	Simplified/grouped route of chemical administration 
Lipodyst	-	Presence of lipodystrophy (Yes/no) 
Obesity	-	Reported presence of obesity (Yes/No/? (unclear)/blank) - left blank if not stated. Yes=statistically greater weight (or fat mass) in model than control animals. 
Insul_res	-	Reported presence of insulin resistance (Yes/No/? (unclear)/blank) - left blank if not stated. Yes=statistically higher fasting/post-prandial insulin or glucose in model than control animals. 
Dyslip	-	Reported presence of dyslipidaemia (Yes/No/? (unclear)/blank) - left blank if not stated. Yes=statistically higher triglycerides/low density lipoprotein/cholesterol or lower high density lipoportein in model than control animals. 
Metab_score	-	Metabolic syndrome score (0-3) from the previous three items 
Liver_score	-	Liver histology score (0-11) from the subsequent 11 times 
NAFLD_score	-	Sum of metabolic and liver scores (0-14) 
Transamin	-	Reported presence of raised transaminases (Yes/No/? (unclear)/blank) - left blank if not stated. Yes=statistically higher alanine or aspartate aminotransferase in model than control animals. 
Balloon_YN	-	Reported presence of hepatocellular ballooning at any time point (Yes/No/? (unclear)/blank) - left blank if not stated 
NASH_YN	-	Reported presence of steatohepatitis at any time point (Yes/No/? (unclear)/blank) - left blank if not stated 
Lobul_YN	-	Reported presence of lobular inflammation at any time point (Yes/No/? (unclear)/blank) - left blank if not stated 
Portal_YN	-	Reported presence of (peri-)portal inflammation at any time point (Yes/No/? (unclear)/blank) - left blank if not stated 
Fib_YN	-	Reported presence of fibrosis of any stage at any time point (Yes/No/? (unclear)/blank) - left blank if not stated 
Fib_F1	-	Reported presence of at least stage 1 fibrosis (Yes/No/? (unclear)/blank) - left blank if not stated 
Fib_F2	-	Reported presence of at least stage 2 fibrosis (Yes/No/? (unclear)/blank) - left blank if not stated 
Fib_F3	-	Reported presence of at least stage 3 fibrosis (Yes/No/? (unclear)/blank) - left blank if not stated 
Fib_F4	-	Reported presence of at least stage 4 fibrosis (Yes/No/? (unclear)/blank) - left blank if not stated 
HCC_YN	-	Reported presence of hepatocellular carcinoma at any time point (Yes/No/? (unclear)/blank) - left blank if not stated 
Fib_stage	-	Maximum fibrosis stage reported for the model (or Yes_NR if stated to show fibrosis but stage not described) 
HepTrig	-	Reported presence of elevated quantitatively measured hepatic triglyceride content (Yes/No/? (unclear)/blank) - left blank if not stated. Yes=statistically higher hepatic triglyceride in model than control animals. 
Steat_YN	-	Reported presence of hepatic steatosis (Yes/No/? (unclear)/blank) - left blank if not stated. 
Steat_wk	-	Timings (in weeks) that steatosis was described in the model 
Steat_max	-	Maximum age (in weeks) that the presence of steatosis was described for this model 
Balloon_wk	-	Timings (in weeks) that ballooning was described in the model 
Balloon_max	-	Maximum age (in weeks) that the presence of ballooning was described for this model 
NASH_wk	-	Timings (in weeks) that steatohepatitis was described in the model 
NASH_max	-	Maximum age (in weeks) that the presence of steatohepatitis was described for this model 
Lobul_wk	-	Timings (in weeks) that lobular inflammation was described in the model 
Lobul_max	-	Maximum age (in weeks) that the presence of lobular inflammation was described for this model 
Portal_wk	-	Timings (in weeks) that portal inflammation was described in the model 
Portal_max	-	Maximum age (in weeks) that the presence of portal inflammation was described for this model 
Fib_wk	-	Timings (in weeks) that fibrosis was described in the model 
Fib_max	-	Maximum age (in weeks) that the presence of fibrosis was described for this model 
HCC_wk	-	Timings (in weeks) that hepatocellular carcinoma was described in the model 
HCC_max	-	Maximum age (in weeks) that the presence of hepatocellular carcinoma was described for this model 
Timing_wk	-	How to interpret the 'timings (weeks)' in this row (model). Usually as number of weeks after diet was started ('Diet') or week of life ('Life'). 
Ref_det	-	The IDs of the individual studies that utilised this model (see column A in tab 1) 
Ref_no	-	The number of studies using this model 
Comments	-	Additional comments 
